# Antigen-derived peptides directly engage the unfolded-protein sensor IRE1α to curb cross-presentation by dendritic cells

**DOI:** 10.1101/2021.09.10.459738

**Authors:** Ofer Guttman, Adrien Le Thomas, Scot Marsters, David A. Lawrence, Lauren Gutgesell, Jonathan M. Harnoss, Simone M. Haag, Aditya Murthy, Geraldine Strasser, Zora Modrusan, Thomas Wu, Ira Mellman, Avi Ashkenazi

## Abstract

Dendritic cells (DCs) promote adaptive immunity by cross-presenting antigen-based epitopes to CD8^+^ T cells. DCs process internalized protein antigens into peptides that enter the endoplasmic reticulum (ER) and upload onto major histocompatibility type I (MHC-I) protein complexes for cell-surface transport and cross-presentation. Perplexingly, DCs often exhibit activation of the ER-stress sensor IRE1α in the absence of classical ER stress—leaving the underlying mechanism unexplained. Here we show that antigen-derived hydrophobic peptides directly engage ER-resident IRE1α by masquerading as unfolded proteins. Furthermore, IRE1α activation depletes MHC-I heavy-chain mRNAs through regulated IRE1α-dependent decay (RIDD), thereby curtailing antigen cross-presentation. In tumor-bearing mice, IRE1α disruption increased MHC-I expression on tumor-infiltrating DCs, and enhanced recruitment and activation of CD8^+^ T cells. Moreover, IRE1α inhibition synergized with anti-PD-L1 antibody treatment to cause tumor regression. Our findings elucidate the mechanism and consequence of antigen-driven IRE1α activation in DCs, yielding a promising combination strategy for cancer immunotherapy.

## INTRODUCTION

Dendritic cells (DCs) comprise a unique myeloid cell subset that plays a crucial role in antigen presentation during the development and elaboration of adaptive immunity (Mellman & Steinman, 2001; Steinman, 2007). Certain DC lineages mediate the specialized process of antigen cross-presentation, which initiates cytotoxic CD8^+^ T cell responses (Gatti & Pierre, 2003; Mellman & Steinman, 2001; Palucka, Banchereau, & Mellman, 2010). DCs are highly proficient in acquiring antigens from tissue microenvironments, through endocytosis of soluble proteins or phagocytosis of cell fragments and corpses (Z. Chen et al., 2001; Sallusto, Cella, Danieli, & Lanzavecchia, 1995). During exposure to a pulse of protein antigen, DCs internalize the polypeptide, which reaches the cytoplasm and undergoes proteasomal processing into shorter peptides (Alloatti, Kotsias, Magalhaes, & Amigorena, 2016). Subsequently, the transporter associated with antigen processing (TAP), which resides in the ER membrane, enables importation of the peptides into the ER lumen. Within the ER, the peptides are uploaded through chaperone-aided events onto MHC-I protein complexes, which are composed of heavy and light polypeptide chains (Jhunjhunwala, Hammer, & Delamarre, 2021; Thomas & Tampe, 2017). The peptide-MHC complexes traffic to the DC surface, where the epitopes are cross-presented to engage cognate T cell receptors (TCRs) on juxtaposing T cells. In cancer, both intrinsic and therapeutic mechanisms require efficient DC-mediated cross-presentation of tumor antigens to CD8^+^ T cells to achieve effective anti-tumor immunity (Barber et al., 2006; Barry et al., 2018; D. S. Chen & Mellman, 2013; Garris et al., 2018; Jansen et al., 2019; Spranger et al., 2016). However, few treatment strategies are currently available to directly modulate DC cross-presentation.

The ER mediates 3D folding of newly synthesized proteins that are destined for plasma membrane insertion or extracellular secretion. Elevated demand for protein folding causes ER stress and triggers the unfolded protein response (UPR), which drives ER adaptation to restore homeostasis (Hetz, 2012; Walter & Ron, 2011; Wang & Kaufman, 2016). The mammalian UPR comprises three key ER-transmembrane proteins: IRE1α, PERK, and ATF6. IRE1α senses ER stress mainly through its ER-lumenal domain, and transmits intracellular signals via a cytoplasmic kinase-endoribonuclease module (Cox, Shamu, & Walter, 1993; K. P. Lee et al., 2008). IRE1α detects misfolded proteins through indirect and direct mechanisms: Indirect engagement involves unfolded-protein binding to the ER chaperone BiP/GRP78, which otherwise keeps IRE1α in check (Amin-Wetzel et al., 2017; Bertolotti, Zhang, Hendershot, Harding, & Ron, 2000). Direct engagement involves unfolded-protein binding to the lumenal domain of IRE1α, through exposed hydrophobic regions that are otherwise buried within correctly folded proteins (Gardner & Walter, 2011; Karagoz et al., 2017). IRE1α activates through molecular events that involve dimerization, kinase trans-autophosphorylation, oligomerization, and consequent endoribonuclease (RNase) engagement (Korennykh et al., 2009; Tirasophon, Welihinda, & Kaufman, 1998).

The IRE1α RNase performs two central functions: (1) activation of the transcription factor X-box protein 1 spliced (XBP1s) (Hetz, 2012; Walter & Ron, 2011; Wang & Kaufman, 2016); (2) depletion of select mRNAs through the process of RIDD (Hollien et al., 2009; Hollien & Weissman, 2006). XBP1s induces multiple genes that support ER-mediated protein folding, as well as ER-associated degradation (ERAD) of misfolded proteins (Acosta-Alvear et al., 2007; Brodsky, 2012; A. H. Lee, Iwakoshi, & Glimcher, 2003; Smith, Ploegh, & Weissman, 2011; Travers et al., 2000). RIDD on the other hand depletes specific ER-targeted mRNAs to abate ER load (Hollien, 2013; Hollien et al., 2009; Hollien & Weissman, 2006; Lhomond et al., 2018; Maurel, Chevet, Tavernier, & Gerlo, 2014). RIDD also regulates additional cellular functions by degrading mRNAs whose products control triglyceride and cholesterol metabolism (So et al., 2012); apoptosis (Chang et al., 2018; Lam, Marsters, Ashkenazi, & Walter, 2020; M. Lu et al., 2014); autophagy (Bae, Moore, Mella, Hayashi, & Hollien, 2019); antibody production (Tang et al., 2018); and DNA repair (Dufey et al., 2020; Tang et al., 2018).

To process XBP1 mRNA for splicing, the IRE1α RNase recognizes two stem-loop structures, located 26 nucleotides apart; each loop contains the consensus sequence endomotif CNGCAGC (Calfon et al., 2002; Shen et al., 2001; Yoshida, Matsui, Yamamoto, Okada, & Mori, 2001). IRE1α cleaves between the guanine (G) at position 3 and the cytosine (C) at position 4 (Hooks & Griffiths-Jones, 2011; Peschek, Acosta-Alvear, Mendez, & Walter, 2015). Subsequently, RtcB ligates the resulting 5’ and 3’ RNA exons are to produce XBP1s (Jurkin et al., 2014; Kosmaczewski et al., 2014; Lu, Liang, & Wang, 2014; Peschek et al., 2015). In mammals, RIDD typically requires an XBP1-like endomotif, CNGCAGN, within a predicted stem-loop structure (Hollien, 2013; K. Moore & Hollien, 2015; K. A. Moore, Plant, Gaddam, Craft, & Hollien, 2013; Oikawa, Tokuda, Hosoda, & Iwawaki, 2010). Experimental XBP1s disruption artificially increases IRE1α autophosphorylation and augments RIDD (X. Chen et al., 2014; M. Lu et al., 2014; Osorio et al., 2014).

DCs exhibit IRE1α activation in the absence of canonical ER stress (Iwakoshi, Pypaert, & Glimcher, 2007; Osorio et al., 2014; Tavernier et al., 2017). Gene knockout (KO) studies of *XBP1* have indicated divergent effects on antigen cross-presentation in different types of DCs. In CD8α^+^ DCs, *XBP1* KO led to a hyper-activated RIDD phenotype, which disrupted T cell activation by depleting mRNAs encoding specific components of the cross-presentation machinery, *i.e.*, Lamp-1 and TAP binding protein (TAPBP) (Iwakoshi et al., 2007; Osorio et al., 2014; Tavernier et al., 2017). In contrast, in tumor-associated DCs, *XBP1* KO led to changes in lipid metabolism, which improved cross-presentation of tumor antigens and consequent anti-tumor T cell activity (Cubillos-Ruiz et al., 2015). In conventional (c) DC1 subpopulations residing in the lung, *XBP1* KO together with partial *IRE1α* gene disruption reduced DC viability (Tavernier et al., 2017). Pulsing of bone marrow-derived DCs (BMDCs) with melanoma cell lysates as a source of antigens upregulated XBP1s without affecting RIDD, while IRE1α inhibition attenuated cross-presentation to CD8^+^ T cells (Medel et al., 2018). While these studies implicate IRE1α in the biological regulation of DCs, the mechanism underpinning the activation of IRE1α in these cells in the absence of canonical ER stress remains a mystery.

In the present study, we reveal that antigen-derived peptides can directly engage IRE1α in antigen-pulsed DCs by mimicking the action of misfolded proteins. We further show that antigen-induced IRE1α stimulation curtails cross-presentation, through specific RIDD-mediated depletion of MHC-I heavy-chain mRNAs. Blocking this negative feedback in tumor-bearing mice by inhibiting IRE1α upregulated MHC-I levels on DCs and enhanced tumor recruitment and activation of CD8^+^ T cells. Moreover, IRE1α inhibition cooperated with anti-PD-L1 immune-checkpoint disruption to cause tumor regression. Our findings elucidate an unexpected mechanism underlying IRE1α activation in DCs, revealing a promising combinatorial strategy for cancer immunotherapy.

## RESULTS

### Antigen pulsing of BMDCs activates IRE1α

To seek insight into the mechanism of IRE1α activation in DCs, we pulsed mouse BMDCs, matured *ex vivo* with GM-CSF plus IL-4, with the classical protein antigen ovalbumin. In order to cross-present ovalbumin, DCs must internalize the pulsed protein and process it intracellularly; in contrast, DCs can directly present the ovalbumin-based octapeptide SIINFEKL, based on its ability to displace antigens already bound to MHC-I complexes at the cell surface (Alloatti et al., 2016). Ovalbumin pulsing of BMDCs induced concentration- and time-dependent activation of IRE1α, evident by increased protein levels of XBP1s and phospho-IRE1α (Figure 1A and 1B). Comparable to the toll-like receptor (TLR) agonist polyinosinic:polycytidylic acid (poly-I:C)—previously shown to activate IRE1α in leukocytes (Martinon, Chen, Lee, & Glimcher, 2010)—ovalbumin pulsing induced much weaker IRE1α activation than did the potent pharmacological ER-stressor tunicamycin (Figure 1B). In contrast to ovalbumin, SIINFEKL had little effect on IRE1α activity (Figure 1B), suggesting a requirement for intracellular events. Unlike IRE1α, other UPR sensors, *i.e.*, PERK (assessed by induction of its downstream targets ATF4 and CHOP) and ATF6 (assessed by its proteolytic processing), which responded to ER-stress induction by the proteasome inhibitor MG132, did not show measurable activation in response to ovalbumin pulsing (Figure S1A), indicating specific IRE1α engagement in the absence of classical ER stress. To verify *bona fide* IRE1α activation during ovalbumin pulsing, we added the highly selective kinase-based small molecule IRE1α inhibitor, G03089668 (G9668) (Figure S1B-D) (Harnoss et al., 2020; Harnoss et al., 2019; Harrington et al., 2015), which completely blocked IRE1α phosphorylation and XBP1s induction. During a 4-hour pulse of ovalbumin, IRE1α phosphorylation peaked at 2 hours while XBP1s protein peaked at 4 hours (Figure 1C and S1A). Of note, a 1-hour pulsing with ovalbumin followed by washing and incubation in standard media for another 3 hours generated similar levels of XBP1s (data not shown). Taken together, these results indicate a rapid, yet transient, stimulation of IRE1α in BMDCs in response to ovalbumin pulsing. BMDC pulsing with lysates derived from CT26, 4T1 and EMT6 tumor cells (see below) also activated IRE1α (Figure 1D). Moreover, pulsing with a different, highly purified recombinant protein, *i.e.*, clinical-grade soluble CD4, also led to IRE1α activation (Figure 1E), indicating a specific stimulation mechanism independent of potential contamination with bacterial endotoxin (lipopolysaccharide, LPS).

**Figure 1.**
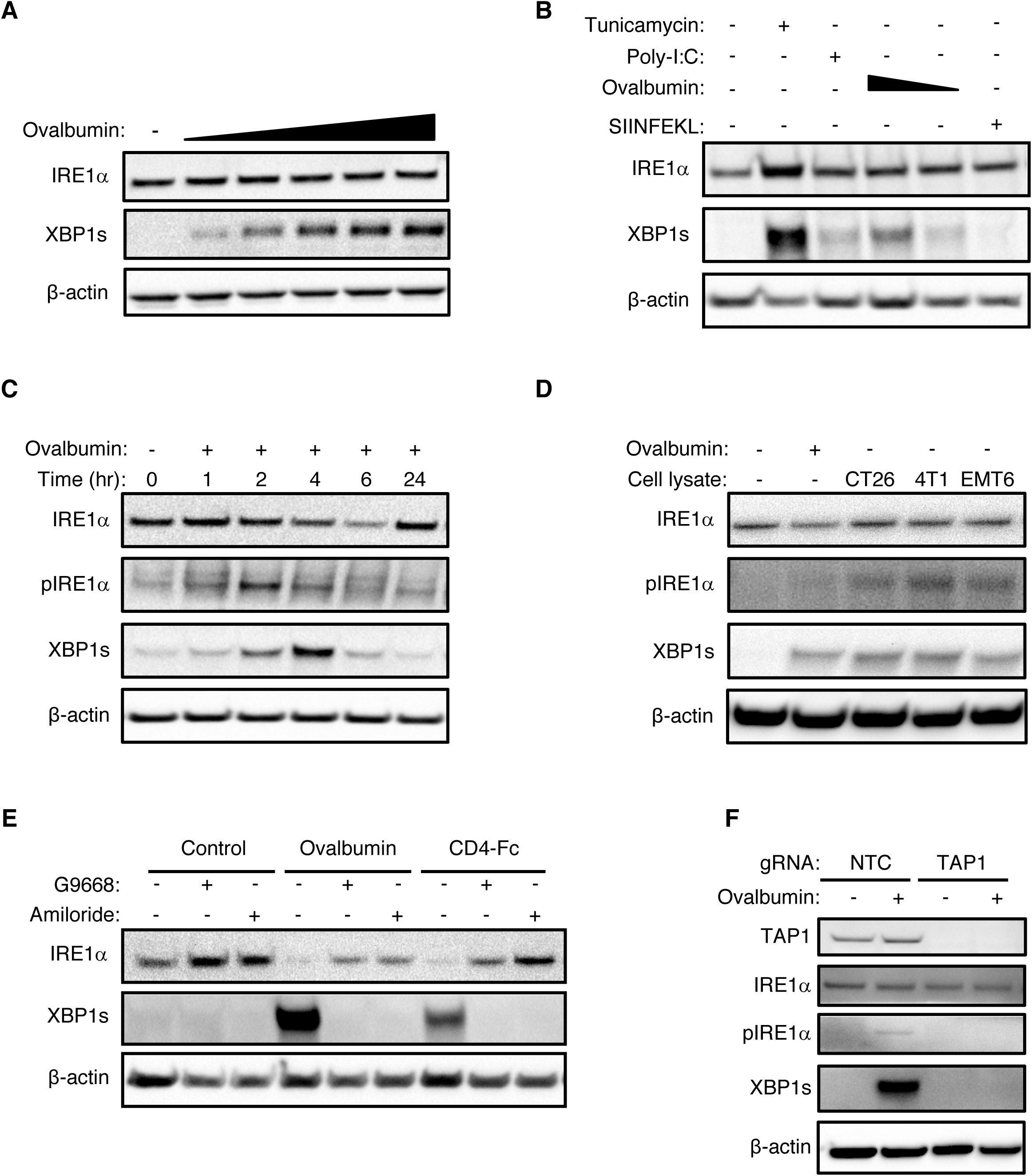
Antigen pulsing of BMDCs activates IRE1α. (**A)** BMDCs were pulsed with ovalbumin (starting at 62.5 μg/ml and sequentially doubled) and analyzed by immunoblot (IB) for the indicated markers. **(B)** BMDCs were pulsed with ovalbumin (starting at 500 μg/ml and sequentially halved) or SIINFEKL (1 μM), or stimulated with tunicamycin (1 μg/ml) or poly-I:C (25 μg/ml) for 4 hr, and analyzed by IB. **(C)** BMDCs were pulsed with ovalbumin (500 μg/ml) for the indicated time and analyzed by IB. **(D)** BMDCs were pulsed for 4 hr with ovalbumin or lysates derived from the indicated cell lines (500 μg/ml protein) and analyzed by IB. **(E)** BMDCs were pulsed for 4 hr with ovalbumin or human soluble CD4-Fc fusion protein (both at 500 μg/ml), combined with DMSO or G9668 (3 μM) or amiloride (10 μM), and analyzed by IB. **(F)** Upon removal from bone marrow, total bone marrow cells were transfected with non-targeting control (NTC) or TAP1-targeting guide RNAs (gRNA), along with CRISPR/Cas9 delivery constructs. Cells were incubated in BMDC-differentiation media for 9 days, pulsed with ovalbumin (500 μg/ml) for 4 hr, and analyzed by IB. All western blot images are representative of at least two similar experiments.

The micropinocytosis inhibitor amiloride (Koivusalo et al., 2010) blocked antigen-induced IRE1α stimulation in BMDCs (Figure 1E and S1E), demonstrating a requirement for antigen internalization. Furthermore, CRISPR/Cas9-based disruption of the *TAP1* gene prevented IRE1α stimulation in response to ovalbumin (Figure 1F), indicating a requirement for antigenic peptide importation into the ER. As expected (Martinon et al., 2010), BMDCs lacking the TLR adapter MyD88 failed to activate IRE1α in response to LPS (Figure S1F); importantly, these cells showed unimpeded IRE1α stimulation in response to ovalbumin, further validating an LPS-independent IRE1α activation. We obtained similar evidence for LPS-independent IRE1α activation upon ovalbumin pulsing of Flt3-ligand (Flt3L)-matured BMDCs from WT or MyD88 KO mice (data not shown). Moreover, unlike BMDCs, mouse embryonic fibroblasts (MEFs) did not display detectable IRE1α activation after exposure to ovalbumin (Figure S1G), suggesting a BMDC-specific mechanism. Taken together, these results suggest that IRE1α engagement in DCs in response to a pulse of a protein antigen occurs independently of LPS and requires antigen uptake as well as importation of antigen-derived peptides into the ER.

### Antigen-derived peptides can directly engage IRE1α

In the context of classical ER stress, otherwise buried hydrophobic segments of unfolded proteins can directly engage the ER-lumenal domain of IRE1α (Gardner & Walter, 2011). Although general UPR activation was absent during antigen pulsing of DCs (Figure S1A and S1B), we reasoned that antigen-derived peptides may directly engage IRE1α by mimicking the action of unfolded proteins. To examine this possibility, we first compared the ability of heat-denatured and native forms of the ovalbumin antigen to bind to a recombinant protein comprising the IRE1α lumenal domain fused to an Fc tag (LD-Fc). Whereas heat-denatured ovalbumin displayed specific and saturable binding to immobilized IRE1α LD-Fc, native ovalbumin showed little binding over background (Figure 2A). We estimated a Kd of ∼ 300 ± 75 μM for denatured ovalbumin, in line with affinities previously reported for unfolded-protein binding to IRE1α LD (Gardner & Walter, 2011). Co-immunoprecipitation studies confirmed concentration-dependent association of heat-denatured ovalbumin to IRE1α LD-Fc, whereas native ovalbumin again did not show appreciable binding (Figure 2B). We obtained similar results by acidic (pH=2) or basic (pH=10) ovalbumin denaturation (data not shown). Thus, ovalbumin denaturation permits direct binding to IRE1α‘s LD.

**Figure 2.**
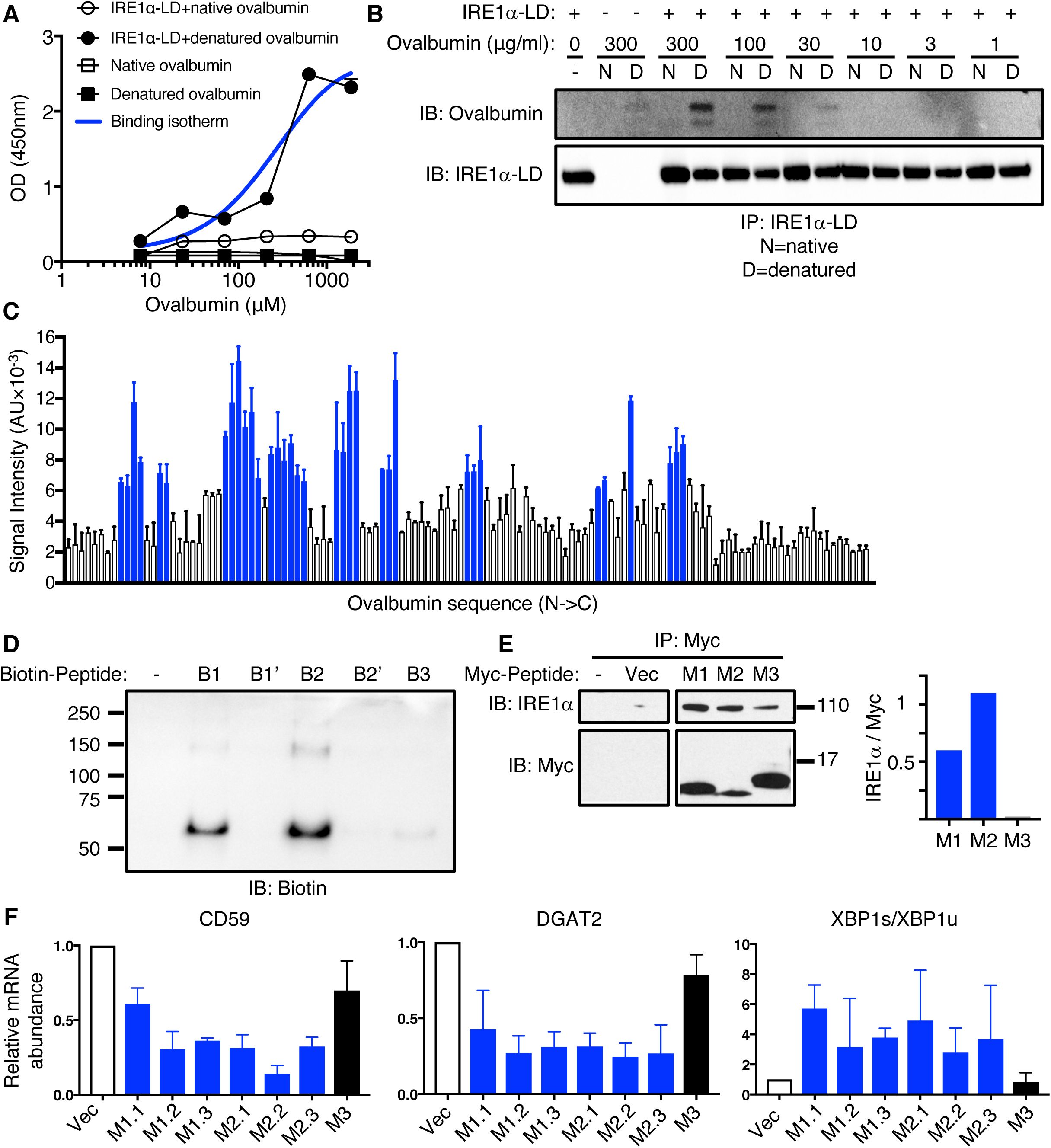
Antigen-derived peptides can directly engage IRE1α. **(A)** Polystyrene wells were coated with native or heat-denatured ovalbumin (10 μg/ml) and incubated with purified recombinant human IRE1α LD-Fc fusion protein (10 μg/ml), followed by colorimetric detection with an HRP-conjugated anti-human Fc antibody. **(B)** Native or heat-denatured ovalbumin at indicated concentrations was incubated with IRE1α LD-Fc (10 μg/ml), immunoprecipitated via monoclonal anti-IRE1α LD antibody, and analyzed by IB. **(C)** A tiled 18 aa-long peptide array spanning ovalbumin was incubated with IRE1α LD-Fc (500 nM) followed by colorimetric detection with an HRP-conjugated anti-human Fc antibody. **(D)** Biotin-tagged ovalbumin-based peptides (100 μM) were incubated with FLAG-tagged IRE1α LD (50 μM), cross-linked with disuccinimidyl suberate (DSS) and analyzed by IB. B1, B2, B3 are WT peptides; B1’, B2’ are mutant peptides in which all hydrophobic residues were replaced by aspartic acid. **(E)** HEK293 cells were transfected with cDNA constructs encoding Myc-tagged peptides (M1, M2, M3) derived from corresponding ovalbumin regions (B1, B2, B3) and containing an ER-directed signal sequence for 48 hr, followed by immunoprecipitation with anti-Myc antibody and IB for IRE1α or Myc. Bar graph indicates signal ratio for IRE1α over Myc. **(F)** U2OS cells were transfected with cDNA constructs encoding Myc-tagged peptides (M1.1, M1.2, M1.3, M2.1, M2.2, M2.3, M3) derived from corresponding ovalbumin regions (B1, B2, B3) and containing an ER-targeting signal sequence for 48 hr, followed by real-time quantitative PCR (RT-qPCR) analysis for XBP1s, XBP1u, CD59, and DGAT2 mRNA levels. Bar graphs in panels **A**, **C** and **F** represent mean ± SD from three independent technical repeats; images in panels **B, D** and **E** represent at least two similar experiments.

Next, to determine whether specific ovalbumin subsegments could interact with IRE1α, we generated a “tiled” peptide array spanning the polypeptide sequence—consisting of 18 amino-acid long synthetic peptides with a 3-residue overlap, as previously described (Gardner & Walter, 2011). We spotted the peptides onto a membrane and examined binding of IRE1α LD-Fc, using horseradish peroxidase-based colorimetric detection. To test an additional antigen, we generated a similar peptide array based on the GP70 protein, which is expressed by CT26 colorectal cancer cells (Takeda et al., 2000). The analyses revealed that 34/123 ovalbumin peptides (clustered in 10 regions) and 64/210 GP70 peptides (19 regions) displayed significant binding to IRE1α LD-Fc (Figure 2C and S2A). In contrast, arrayed peptides derived from these antigens did not exhibit detectable binding to CD4-Fc under identical conditions (data not shown).

We next evaluated the importance of hydrophobic side-chains for peptide binding. We synthesized biotin-tagged peptides corresponding to two binding and one non-binding segments of the tiled ovalbumin array (B1, B2, and B3), and mutated variants of the binders with hydrophobic residues substituted by aspartic acids (B1’, B2’) (Figure S2B). We incubated each peptide with FLAG-tagged IRE1α LD, stabilized formed complexes by chemical crosslinking, and visualized them by anti-biotin immunoblotting. Whereas peptide B3 showed no significant interaction, B1 and B2 exhibited specific binding, associating not only with monomers but also with apparent dimers or oligomers of IRE1α LD-FLAG (Figure 2D). In contrast to B1 and B2, mutated peptides B1’ and B2’ failed to show significant binding, indicating a critical role for hydrophobic residues in B1 and B2 for interaction with IRE1α LD.

To test whether ovalbumin-based peptides can associate with IRE1α in a cellular setting, we transfected HEK293 cells with cDNA constructs encoding shorter Myc-tagged versions of B1, B2 and B3 (M1, M2 and M3)—fused to a signal sequence for direct ER targeting (Figure S2B). Immunoprecipitation with anti-Myc antibody followed by immunoblotting with anti-IRE1α revealed specific co-immunoprecipitation of Myc-tagged peptides with IRE1α (Figure 2E). Analyzing the signal ratios for IRE1α over Myc confirmed markedly greater IRE1α interaction for M1 and M2 as compared to M3. To assess functional IRE1α engagement, we transfected U20S cells with similar cDNA plasmids encoding a series of three peptides each representing M1, M2, or M3 (Figure S2B). We measured IRE1α activation by RT-QPCR analysis of mRNA transcripts for XBP1s and XBP1u, as well as the RIDD targets CD59 and DGAT2. The M1 and M2 peptides functionally engaged IRE1α, as evident by upregulation of XBP1s and depletion of the RIDD-targeted mRNAs encoding CD59 and DGAT2 in comparison to non-transfected controls and peptide M3, which showed no activation above vector controls (Figure 2F). Thus, congruent with the results obtained with the corresponding synthetic peptides in a cell-free setting (Figure 2D), ovalbumin-derived, ER-targeted peptides can specifically bind to the ER-resident IRE1α protein within cells and stimulate its endoribonuclease activity in a manner that corresponds to their direct ability to bind to IRE1α.

### IRE1**α** inhibition in BMDCs augments cross-presentation

To examine whether antigen-induced IRE1α activation impacts cross-presentation, we pulsed BMDCs with ovalbumin in the absence or presence of G9668. We then tested BMDC capacity to activate mouse splenic OT-I CD8^+^ T cells, which express a transgenic TCR specific to the SIINFEKL epitope. During SIINFEKL pulsing, IRE1α inhibition with G9668 had little effect; however, upon ovalbumin pulsing, it significantly and reproducibly augmented subsequent induction of OT-I CD8^+^ T-cell proliferation by ∼ 20% (Figure 3A and S3A). To confirm LPS-independent augmentation, we pulsed BMDCs with endotoxin-free ovalbumin (EF-OVA); comparably, IRE1α inhibition in this setting augmented OT-I CD8^+^ T-cell proliferation by ∼ 29% (Figure 3A). Furthermore, G9668 treatment during pulsing of Flt3L-matured BMDCs with either ovalbumin or EF-OVA increased OT-I CD8^+^ T-cell proliferation by 23% or 24%, respectively (Figure 3B and S3A). Importantly, G9668 did not directly affect proliferation or activation of naïve CD4^+^ or CD8^+^ splenic T cells upon TCR stimulation with anti-CD3 plus anti-CD28 antibodies (Figure S3B and S3C). Moreover, in contrast to CD8^+^ T cell stimulation, IRE1α inhibition during ovalbumin pulsing did not alter MHC class II-restricted activation of splenic OT-II transgenic CD4^+^ T cells, which also harbor an ovalbumin-specific TCR transgene (Figure S3D). Taken together, these results indicate that protein-antigen pulsing of BMDCs leads to LPS-independent IRE1α activation, which in turn specifically dampens MHC-I-restricted antigen cross-presentation to CD8^+^ T cells. Functional inhibition of IRE1α reverses this curbing mechanism, enhancing antigen cross-presentation.

**Figure 3.**
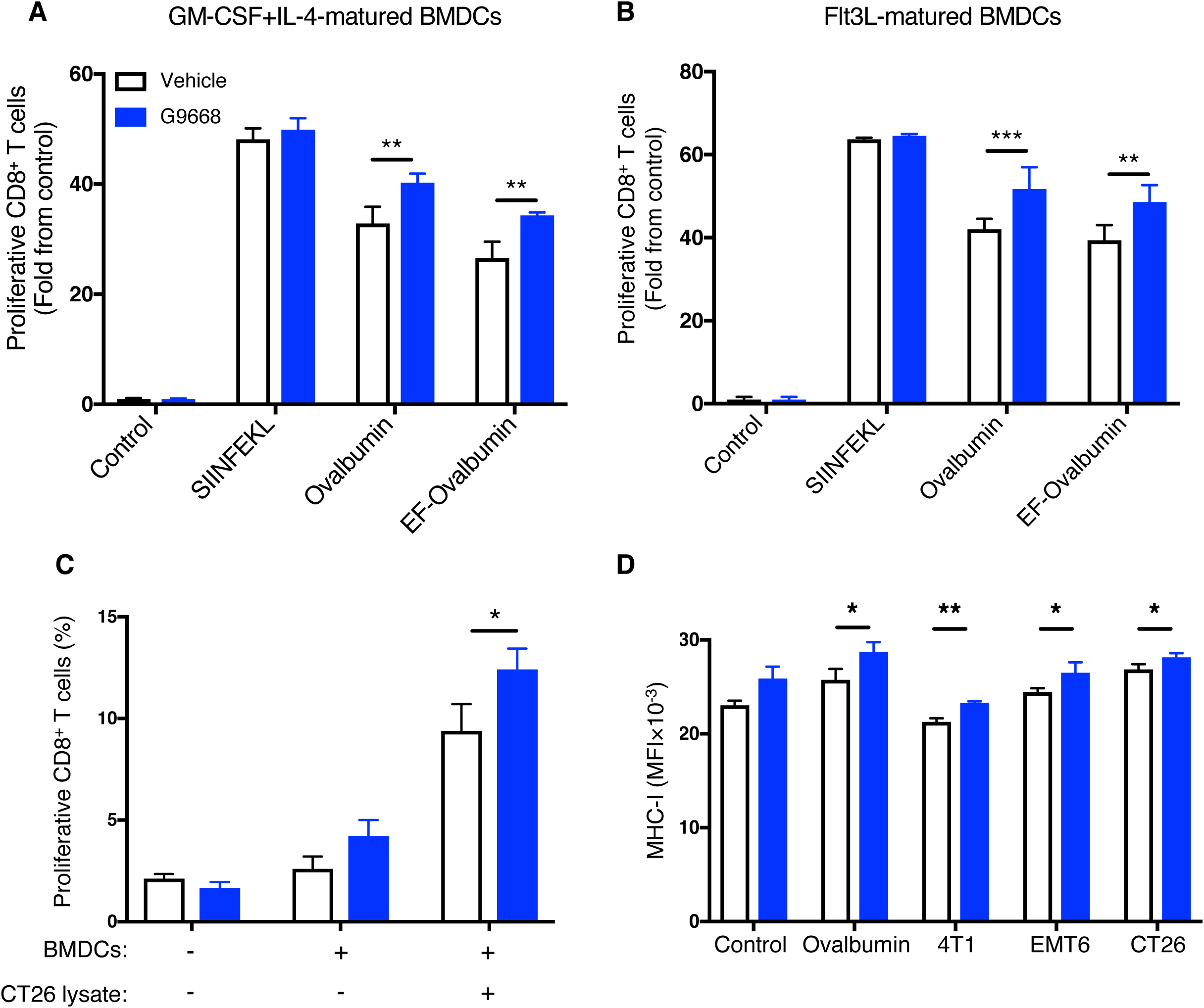
IRE1α inhibition in BMDCs augments antigen cross-presentation to CD8^+^ T cells. **(A, B)** GM-CSF+IL-4-matured **(A)** or Flt3L-matured **(B)** BMDCs were pulsed with ovalbumin (500 μg/ml) with or without G9668 (3 μM) for 24 hr and subsequently co-cultured with magnetically-separated CD8^+^ OT-I T cells for 72 hr, followed by flow cytometry analysis of T cell proliferation by Celltrace Violet. **(C)** BMDCs were pulsed with lysates derived from CT26 cells (500 μg/ml protein) for 24 hr and co-cultured with magnetically-separated CD8^+^ splenic T cells from CT26 tumor-bearing mice for 72 hr, followed by analysis of T cell proliferation by flow cytometry of Celltrace Violet staining. **(D)** BMDCs were pulsed with ovalbumin or lysates derived from 4T1, EMT6 and CT26 cells (500 μg/ml protein) for 8 hr and surface levels of MHC-I were assayed by flow cytometry. Analysis was performed using unpaired, two-tailed *t* test, * P ≤ 0.05 **P ≤ 0.01, ***P ≤ 0.001. Bar graphs in all panels represent mean ± SD from three independent biological repeats; panels **A** and **B** represent data from at least independent experiments collated as fold from control for each experiment.

To investigate the impact of IRE1α inhibition on cross-presentation of tumor antigens, we subcutaneously inoculated BALB/c mice with CT26 cells and allowed tumors to form. We then isolated splenic CD8^+^ T cells (likely possessing TCRs that can recognize CT26 antigens) from these mice and co-incubated them with BMDCs pre-pulsed with CT26 cell lysates. Addition of G9668 augmented cross-presentation of CT26 epitopes to cognate splenic CD8^+^ T cells by ∼ 25% (Figure 3C), in keeping with ovalbumin cross-presentation. Thus, IRE1α inhibition in BMDCs enhances cross-presentation of tumor-derived antigens.

Next, we turned to interrogate which specific aspect of the cross-presentation process is modulated by IRE1α. IRE1α inhibition did not alter the uptake of fluorescently-labeled ovalbumin by BMDCs (Figure S3E). On the other hand, IRE1α inhibition during exposure of BMDCs to ovalbumin or cell lysates significantly increased MHC-I protein expression at the cell surface (Figure 3D), suggesting negative regulation of antigen-driven MHC-I dynamics by IRE1α.

### IRE1**α** activation depletes MHC-I heavy-chain mRNAs via RIDD

Earlier work shows that in lymph node-resident CD8^+^ DCs RIDD constitutively suppresses mRNA transcripts encoding certain components of the cross-presentation machinery, such as TAPBP (Osorio et al., 2014). In ovalbumin-pulsed BMDCs, IRE1α inhibition minimally impacted mRNA levels of TAPBP, nor did it affect transcript abundance of the MHC-I light-chain β-2 microglobulin (β-2M), TAP1, or ER aminopeptidase ERAP1; however, it markedly upregulated MHC-I H-2K heavy-chain mRNA levels by 2.8-fold (Figure S4A).

We therefore considered the possibility that IRE1α activation in response to antigen pulsing attenuates cross-presentation by decreasing MHC-I heavy-chain mRNAs through RIDD. Supporting this hypothesis, computational examination of mRNA sequences encoding the murine H-2K, H-2D, and H-2L heavy-chains revealed the presence of consensus IRE1α-targeted stem-loop endomotifs (Figure 4A), whereas β-2M mRNA did not contain such sequences. To examine whether IRE1α can directly cleave heavy-chain mRNAs, we incubated a phosphorylated recombinant protein comprising the cytoplasmic kinase-endoribonuclease module of IRE1α with RNA transcripts encoding H-2K, H-2D, and H-2L. IRE1α efficiently cleaved all three RNAs (Figure 4B), supporting the involvement of their RIDD-mediated degradation in curbing cross-presentation.

**Figure 4.**
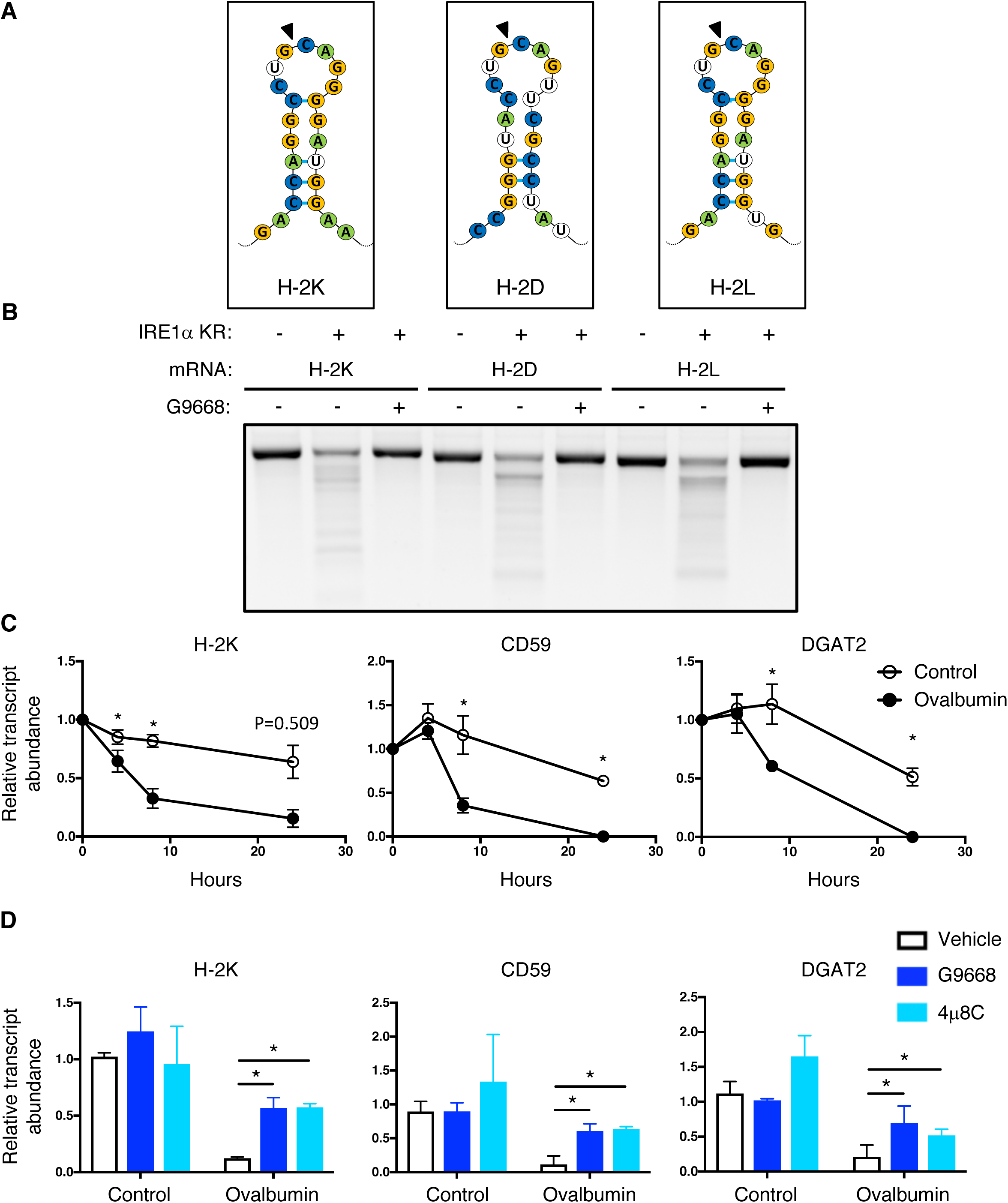
IRE1α activation depletes MHC-I heavy-chain mRNAs via RIDD. **(A)** Consensus stem-loop endomotifs for RIDD recognition found in murine MHC-I heavy-chain H-2K, H-2D and H-2L mRNA sequences. **(B)** Purified recombinant IRE1α kinase-endoribonucleae (KR) protein was incubated with RNA transcripts of H-2K, H-2D and H-2L, in absence or presence of G9668 (10 μM), followed by agarose gel electrophoresis to determine transcript integrity. **(C, D)** BMDCs were treated with actinomycin D (2 μg/ml) to block *de novo* transcription and pulsed with ovalbumin (500 μg/ml) combined with DMSO, G9668 (3 μM), or 4μ8C (1 μM) for the indicated time period **(C)** or for 8 hr **(D)**, followed by RT-qPCR analysis of the indicated transcripts. Analysis was performed using unpaired, two-tailed *t* test, * P ≤ 0.05. Panel **B** image represents three similar experiments; bar graphs in panels **C** and **D** represent mean ± SD from three independent biological repeats.

To examine RIDD-mediated depletion in BMDCs, we prevented *de novo* transcription with actinomycin D. In control-pulsed BMDCs, the mRNA levels of H-2K as well as the canonical RIDD targets CD59 and DGAT2 remained stable over a 24-hour period; in contrast, in antigen-pulsed BMDCs, the levels of both H-2K and CD59 mRNAs substantially declined over time (Figure 4C). Furthermore, both the kinase-based IRE1α inhibitor G9668 and the RNase-directed IRE1α inhibitor 4μ8c (Cross et al., 2012) substantially rescued mRNAs encoding H-2K, CD59, and DGAT2 (Figure 4D), as well as several other known RIDD targets, *i.e.,* BLOS1, RNF213 and IRF7 (Figure S4B), demonstrating RIDD-based depletion.

Thus, antigen pulsing of DCs activates IRE1α, which curbs cross-presentation by depleting MHC-I heavy-chain mRNAs through RIDD.

### IRE1α inhibition in tumor-bearing mice upregulates MHC-I on tumor DCs and augments CD8^+^ T cell engagement

To determine whether the dampening effect of IRE1α on antigen cross-presentation has functional consequences *in vivo*, we examined the impact of IRE1α disruption on syngeneic tumor growth in mice. Tumor-cell-autonomous knockout of *IRE1α* by CRISPR/Cas9 had little effect on subcutaneous growth of CT26 colon tumors (Figure S5A and S5B). In contrast, systemic pharmacological inhibition of IRE1α with G9668 substantially attenuated tumor progression as compared to vehicle treatment (Figure 5A, S5C, and S5D), suggesting enhancement of host-mediated anti-tumor activity. To specifically elucidate potential immune effects, we analyzed by flow cytometry the tumor-associated leukocyte populations after 7 days of treatment. As compared to controls, tumors from G9668-treated mice showed significantly greater infiltration by CD11c^+^ MHC class II^high^ DCs (Figure 5B), specifically belonging to the cDC1 (XCR1^+^ CD103^+^) subpopulation (Figure 5C). Furthermore, tumor-infiltrating cDC1s showed significantly higher surface levels of MHC-I, as well as the RIDD marker CD59, in G9668-treated mice as compared to controls (Figure 5D). Moreover, G9668 treatment led to significantly greater numbers of tumor-infiltrating cytotoxic CD8^+^ T cells (Figure 5E), and to higher expression by these cells of the activation markers granzyme B, PD-1, and CD44 (Figure 5F). Importantly, staining with recombinant MHC-I tetramer complexes presenting the CT26 tumor antigen GP70 revealed significantly higher levels of tumor-infiltrating GP70-specific CD8^+^ T cells in G9668-treated mice (Figure 5G). Thus, IRE1α inhibition attenuates CT26 tumor growth in conjunction with elevated tumor infiltration, MHC-I expression, and tumor-antigen cross-presentation by cDC1s, augmenting the recruitment and activation of tumor-reactive CD8^+^ T cells.

**Figure 5.**
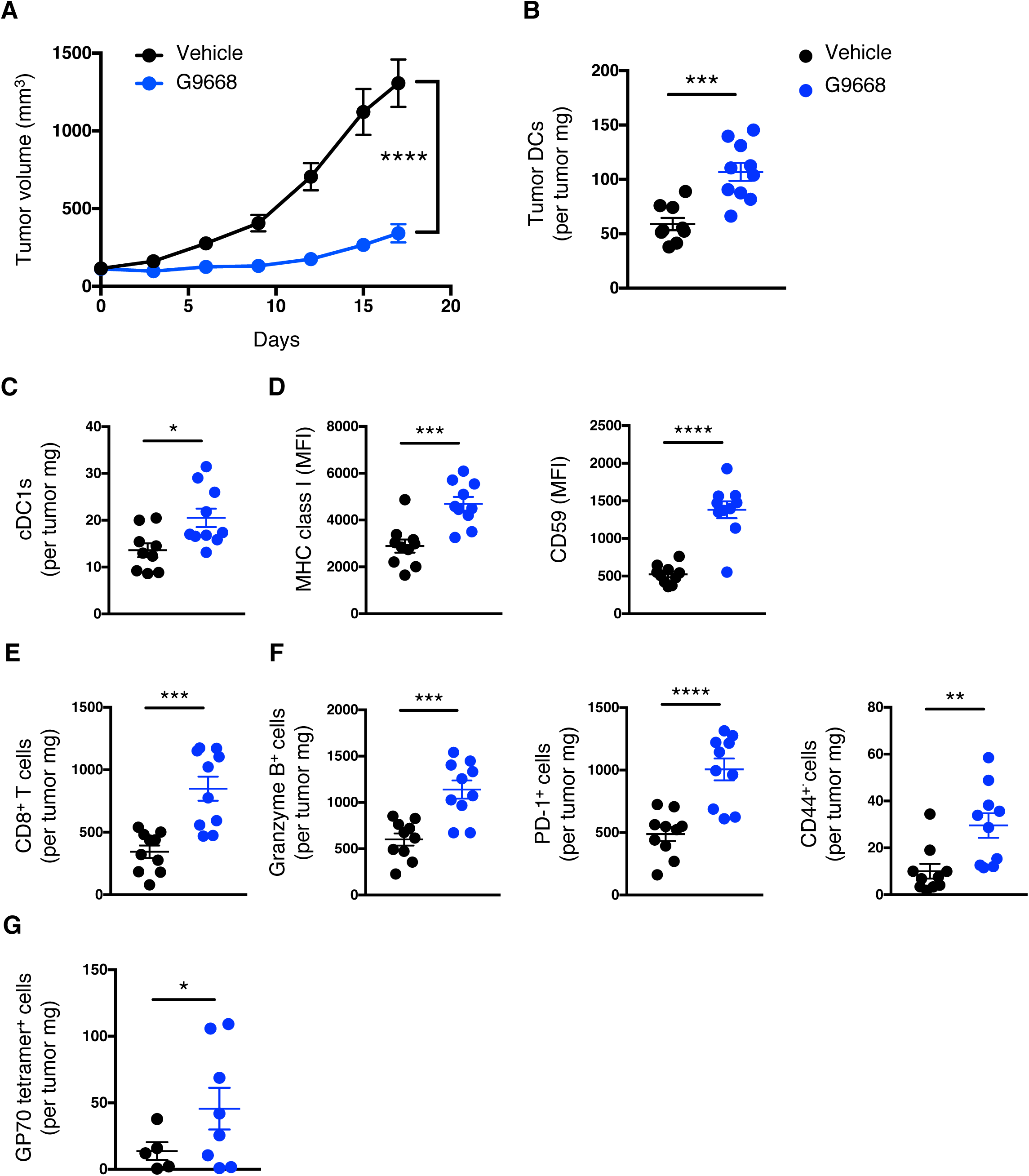
IRE1α inhibition attenuates CT26 tumor growth in conjunction with enhanced MHC-I expression on tumor DCs and CD8^+^ T cell recruitment and activation. Mice were inoculated s.c. with CT26 cells, grouped out 7 days afterwards and treated with vehicle or G9668 (250 mg/kg, BID). **(A)** Growth trajectories of CT26 tumors in vehicle- and G9668-treated animals over 17 days (n = 15). **(B-G)** Flow cytometry analysis of tumor DCs and T cells from mice treated for 7 days. **(B-D)** Quantification of tumor-infiltrating total DCs **(B)** and cDC1s **(C)**; and characterization of cDC1 expression of MHC-I and CD59 **(D)** by flow cytometry (n = 9 for vehicle-treated group and 10 for G9668-treate group). Total DCs were characterized as F4/80^low^, CD11c^+^ and class II MHC^high^, while cDC1s were characterized as CD103^+^ XCR1^+^ CD11b^-^. **(E-F)** Measurement of tumor-infiltrating CD8^+^ T cell abundance **(E)**, expression of indicated activation markers **(F)** (n = 9 for vehicle-treated group and 10 for G9668-treate group), and binding of GP70 tetramers **(G)** (n = 6 for vehicle and 8 for G9668 group). Analysis was performed using one-way ANOVA for panel **A** and unpaired, two-tailed *t* test for panels **B**-**G**, * P ≤ 0.05 *P ≤ 0.05, **P ≤ 0.01, ***P ≤ 0.001, ****P ≤ 0.0001. Scatter plots in all panels represent mean ± SD.

### Single-cell RNAeq demonstrates IRE1α regulation of MHC-I mRNAs in tumor DCs

To examine immune modulation in a different tumor model, we used syngeneic 4T1 triple-negative breast cancer (TNBC) cells. In contrast to the CT26 model (Figure S5A and S5B), cell-autonomous knockout of *IRE1α* in 4T1 cells caused notable tumor-growth inhibition (TGI) of 53% (Figure S6A and S6B). Nevertheless, systemic treatment of mice bearing parental *IRE1α* wildtype 4T1 tumors with G9668 led to a stronger TGI of 82% (Figure 6A, S6C and S6D), suggesting both cell-autonomous and host-mediated anti-tumor effects. The myeloid compartment in tumors has been systematically studied by single-cell RNA sequencing (scRNAseq) (Cheng et al., 2021; Mariathasan et al., 2018). We performed scRNAseq after 6 days of treatment to analyze the tumor leukocytic populations. Tumors in G9668-treated mice showed enrichment in both DCs and CD8^+^ T effector cells, but not in naïve T cells (Figure 6B and 6C). Importantly, tumor-infiltrating DCs in G9668-treated mice displayed significantly higher mRNA levels of H-2K and H-2D heavy-chains; though not of TAPBP transcripts (Figure 6D). CD59 and DGAT2 mRNAs were insufficiently abundant to enable accurate quantification, but five other RIDD targets that were detected, *i.e.* BLOS1, FERMT3, IRF7, RNF213 and SPON1, showed significant increases (Figure S6E), confirming RIDD inhibition. Tumors in G9668-treated mice had unaltered levels of M1 macrophages, but showed significantly fewer M2 macrophages and monocytes as compared to controls (Figure S6F). Flow cytometric analysis after 6 days of treatment showed that IRE1α inhibition significantly increased tumor infiltration by cytotoxic CD8^+^ T cells and their expression of the activation markers IFN-γ, PD-1, and CD69, independent of *IRE1α* status in the malignant cells (Figure 6E). These results further support the conclusion that IRE1α inhibition augments anti-tumor immune responsiveness by increasing MHC-I expression on tumor-associated DCs and consequent engagement of tumor-infiltrating cytotoxic CD8^+^ T cells.

**Figure 6.**
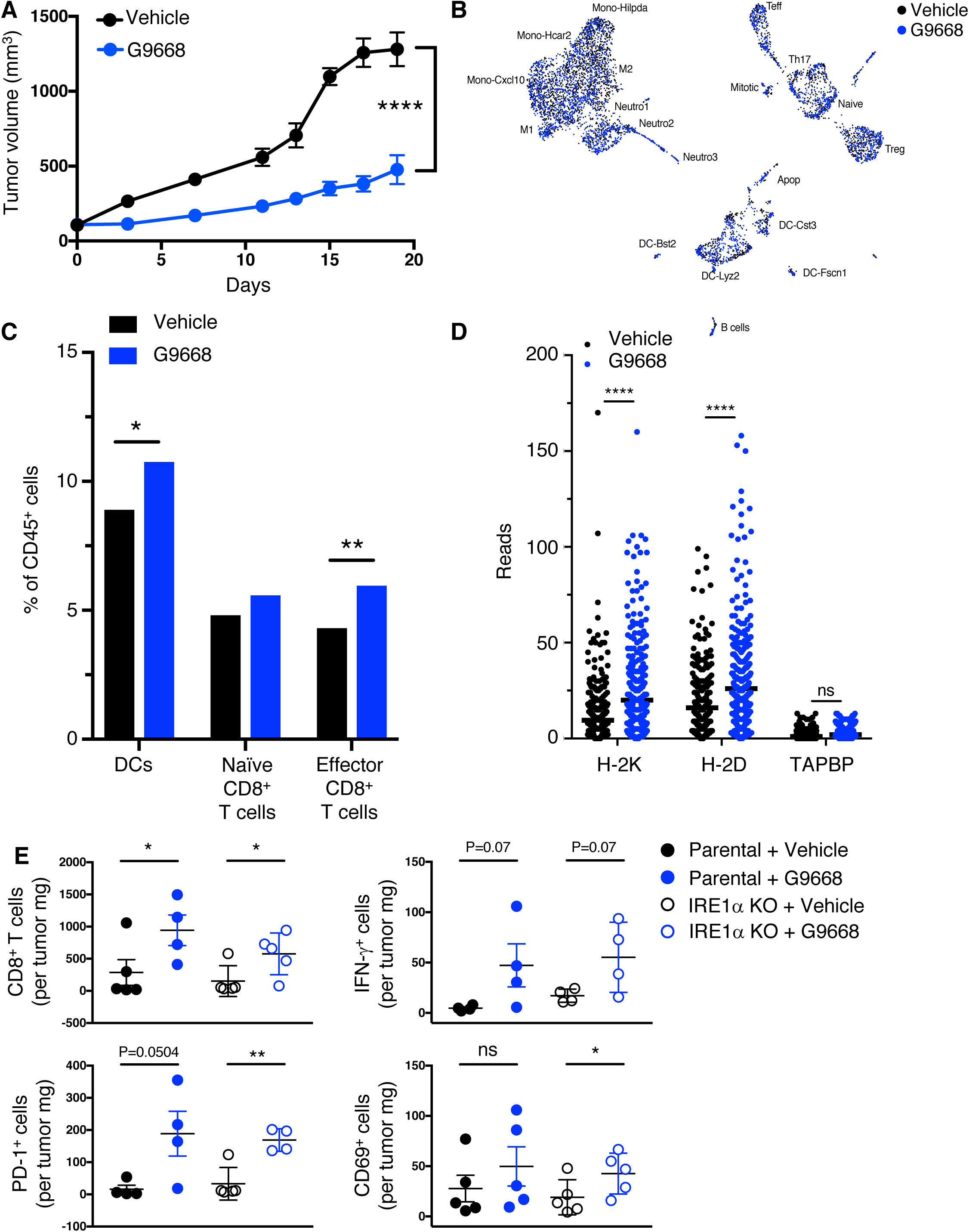
IRE1α inhibition attenuates 4T1 tumor growth in conjunction with increased myeloid MHC-I mRNA levels and DC and CD8^+^ T cell tumor infiltration. **(A-E)** Mice were inoculated s.c. with 4T1 cells, grouped out 7 days afterwards and treated with vehicle or G9668 (250 mg/kg, BID). **(A)** Tumor growth trajectories were measured over 19 days (n = 15). **(B-E)** Mice were treated with G9668 for 6 days and tumor-infiltrating leukocytes were then analyzed. **(B)** Representation of tumor-infiltrating leukocytes by t-distributed stochastic neighbor embedding (t-SNE) plot from single-cell RNA sequencing data. **(C)** Abundance of tumor-infiltrating DCs and CD8^+^ T cells in vehicle- and G9668-treated animals. **(D)** Transcript levels of indicated genes in tumor-infiltrating DCs. **(E)** Mice were inoculated s.c. with parental or IRE1α KO 4T1 cells, grouped out 7 days afterwards and treated with vehicle or G9668 (250 mg/kg) (n = 5). Abundance of CD8^+^ T cells in the tumor and expression of indicated activation markers were assayed by flow cytometry. Analysis was performed using one-way ANOVA for panel **A** and unpaired, two-tailed *t* test for panels **C**-**E**, *P ≤ 0.05, **P ≤ 0.01, ****P ≤ 0.0001. Scatter plots represent mean ± SD.

### Systemic IRE1α inhibition cooperates with immune checkpoint blockade

Although immune-checkpoint disruption has transformed patient benefit in a number of cancer settings (Marinelli et al., 2020), further advances are needed to achieve wider effectiveness. To investigate whether IRE1α inhibition would complement immune-checkpoint blockade, we turned to the orthotopic EMT6 TNBC model, previously found to exhibit partial responsiveness to anti-programmed death ligand (PD-L)1 antibody therapy upon implantation in the mouse mammary fat pad (Mariathasan et al., 2018). Similar to the CT26 model, cell-autonomous *IRE1α* KO in EMT6 cells had minimal impact on tumor growth (Figure S7A and S7B), identifying an additional suitable model to specifically interrogate the impact of IRE1α inhibition on immune modulation. Treatment of EMT6 tumor bearing mice with either the anti-mouse PD-L1 monoclonal antibody 6E11 or with G9668 partially impaired EMT6 tumor progression (Figure 7A and S7C). Remarkably, combined administration of 6E11 and G9668 led to frank tumor regression with a mean TGI rate of 114%, resulting in significantly better efficacy than either monotherapy (p < 0.01). In keeping with the other models, treatment of EMT6 tumor-bearing mice with G9668 for 7 days significantly increased surface levels of MHC-I and CD59 in tumor-infiltrating cDC1s (Figure 7B). Moreover, G9668 increased tumor invasion by cytotoxic CD8^+^ T cells and their expression of the activation markers IFN-γ and granzyme B (Figure 7C). Thus, IRE1α inhibition effectively complements PD-L1-based immune-checkpoint disruption to reverse syngeneic orthotopic tumor progression.

**Figure 7.**
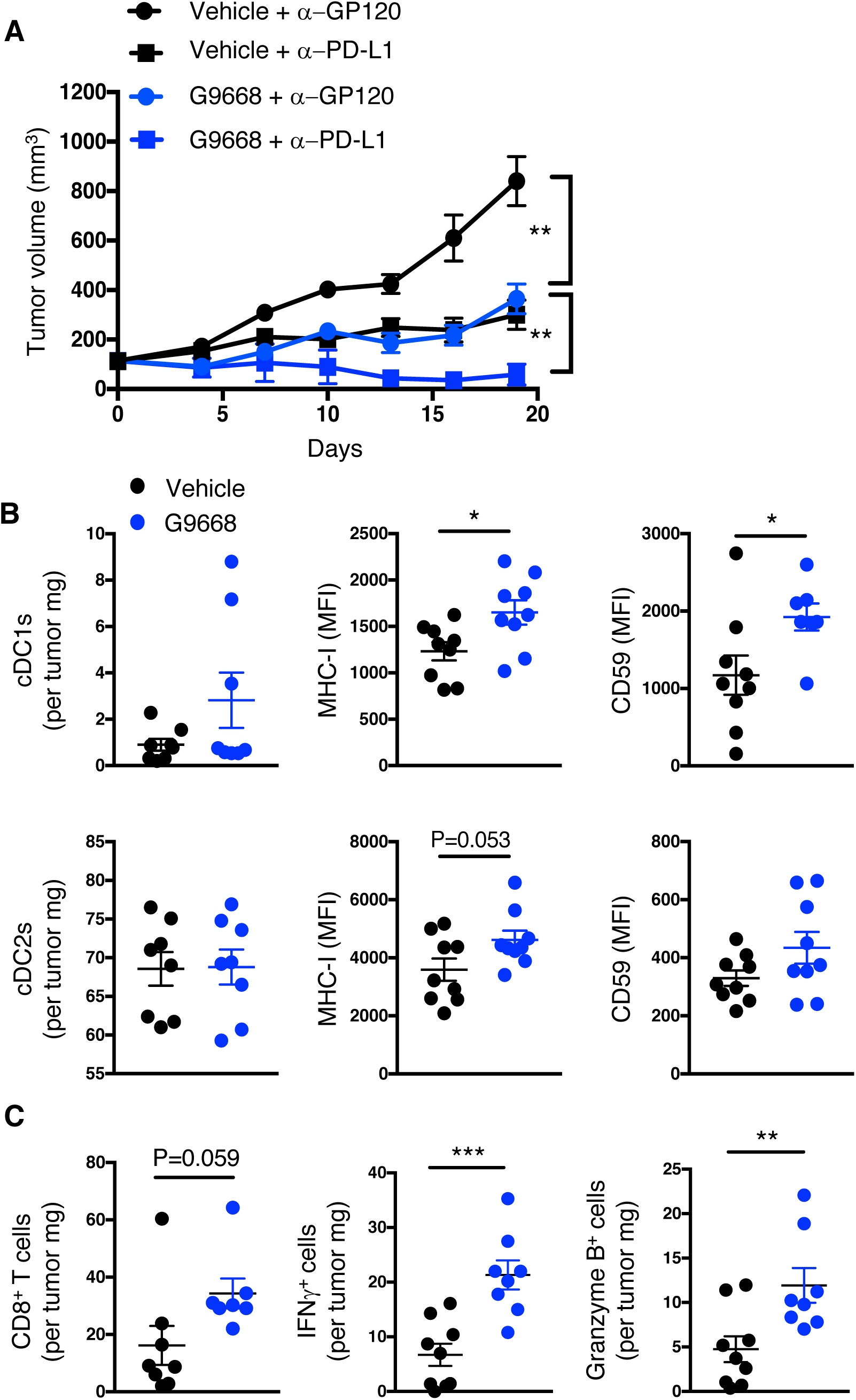
IRE1α inhibition attenuates EMT6 tumor growth and synergizes with anti-PD-L1 antibody in conjunction with increased DC and CD8^+^ T cell tumor infiltration and activation. Mice were inoculated orthotopically with EMT6 cells in the mammary fat pad, grouped out 7 days afterwards and treated with vehicle, G9668 (250 mg/kg, BID), anti-PD-L1 antibody (10 mg/kg at first dose, 5 mg/kg BIW thereafter), or the combination. **(A)** Tumor growth trajectories were measured over 19 days (n = 15). **(B, C)** Mice were treated with G9668 for 7 days and then sacrificed (n = 9). **(B)** Abundance of tumor cDC1s and cDC2s, as well as expression of class I MHC and CD59, were measured by flow cytometry. Total DCs were characterized as F4/80^low^, CD11c^+^ and class II MHC^high^, while cDC1s were characterized as CD103^+^ XCR1^+^ CD11b^-^. **(C)** Abundance and activation marker expression of tumor-infiltrating CD8^+^ T cells were assayed by flow cytometry. Analysis was performed using one-way ANOVA for panel **A** and unpaired, two-tailed *t* test for panels **B** and **C**, * P ≤ 0.05, **P ≤ 0.01, ***P ≤ 0.001. Scatter plots in all panels represent mean ± SD.

## DISCUSSION

The mechanism of IRE1α activation in the absence of classical ER stress in DCs has been mysterious (Iwakoshi et al., 2007; Medel et al., 2018; Osorio et al., 2014; Tavernier et al., 2017). Our present findings reveal that exposure of DCs to pulsed protein antigens drives IRE1α activation through an LPS-independent process analogous to the direct engagement of IRE1α by unfolded proteins under canonical ER stress. We further show that antigen-induced IRE1α activity curbs cross-presentation through RIDD-mediated depletion of MHC-I heavy-chain mRNAs. This functional consequence likely represents a negative feedback loop that fine-tunes cross-presentation, perhaps to prevent inappropriate or excessive T-cell activation during sterile tissue injury. Importantly, the disruption of this inherent negative feedback by inhibiting IRE1α cooperates with immune-checkpoint blockade to enhance anti-tumor immune responses, revealing exciting potential for therapeutic translation.

Our BMDC studies showed that antigen pulsing selectively activates IRE1α but not PERK or ATF6, excluding the standard UPR as a key mechanistic driver. This contrasts with plasmacytoid DCs (pDCs), which do not mediate cross-presentation, and interestingly display constitutive activation of PERK (Mendes et al., 2021). BMDC pulsing with different protein antigens, or with lysates of several cancer cell lines, induced significant levels of IRE1α activity. Although these levels were markedly weaker than those induced by strong pharmacological ER stressors, antigenic IRE1α stimulation was highly reproducible, reaching peak intensity within 2 – 4 hours of exposure, and then declining. These rapid yet transient kinetics are consistent with the time frame of antigen uptake, processing, and ER entry. Indeed, our further mechanistic dissection indicated that antigen-driven IRE1α activation requires both pinocytosis and ER importation events, but not TLR signaling.

IRE1α resides in the ER membrane and responds through its lumenal domain to ER accumulation of unfolded or misfolded proteins during canonical ER stress (Hetz, 2012; Walter & Ron, 2011). Based on our observation that antigen-induced IRE1α activation required TAP1—a critical mediator for importation of antigen-derived peptides into the ER—we reasoned that antigen-based peptides entering the ER may directly engage IRE1α by masquerading as unfolded or misfolded proteins. Several lines of evidence supported this idea. First, whereas native ovalbumin failed to interact with the IRE1α LD, unfolded ovalbumin—generated by denaturation with heat or extreme pH— was capable of direct LD binding, with affinity comparable to that of unfolded proteins (Gardner & Walter, 2011). Second, specific ovalbumin-based peptides bound to the IRE1α LD in a manner that required their hydrophobic amino acids. Third, cellular expression of ER-directed ovalbumin-based peptides demonstrated congruent interaction with, and functional engagement of, cellular IRE1α. The lack of stimulation of the other UPR branches in response to antigen pulsing further indicates a direct IRE1α activation mechanism independent of BiP. Hence, although a nascent protein that is incorrectly folded by the ER and a peptide that lacks 3D structure due to proteolytic cleavage of its parent protein in the cytoplasm are distinct, both can be sensed by IRE1α. Although any protein-producing cell may harbor some constitutive level of IRE1α-peptide interactions, our data suggest that DCs are uniquely capable of IRE1α activation upon transient exposure to a high concentration of an extracellular protein. Moreover, for DCs, which are highly specialized in antigen cross-presentation, such IRE1α activation can have unique functional consequences. The recent discovery of pervasive functional peptide translation in cells (J. Chen et al., 2020) raises an intriguing question of whether additional peptide modalities besides antigen processing may similarly engage IRE1α.

Earlier work interrogating the involvement of IRE1α in DC regulation relied primarily on *XBP1* KO—a strategy that does not completely disrupt, and in some cases even augments, the kinase-endoribonuclease activity of IRE1α. In CD8^+^ DCs, *XBP1* KO causes artificial RIDD hyper-activation, which leads to mRNA depletion of certain components of the cross-presentation machinery, *i.e.*, LAMP-1, TAPBP, and β2M (Osorio et al., 2014). In contrast, BMDC pulsing with melanoma cell lysates activated XBP1 splicing but not RIDD, and XBP1s appeared to promote, rather than disrupt, efficient melanoma antigen cross-presentation (Medel et al., 2018). Direct infection of BMDCs by *Toxoplasma gondii* led to IRE1α activation via MyD88-dependent TLR signaling, with decreased cross-presentation upon *XBP1* KO and partial disruption of IRE1α (Poncet et al., 2021). The partial disruption of IRE1α signaling by *XBP1* KO left the biological consequence of IRE1α engagement during antigen cross-presentation incompletely understood.

To impede IRE1α‘s enzymatic activity more fully, we used the highly selective and potent kinase-based IRE1α inhibitor G9668, which blocks both the kinase and the kinase-controlled endoribonuclease activities of IRE1α. Indeed, G9668 fully disrupted antigen-induced IRE1α activation in DCs, preventing IRE1α auto-phosphorylation, as well as consequent RNase-dependent XBP1 splicing and RIDD. While G9668 did not alter DC-surface presentation of ovalbumin-derived SIINFEKL peptide to OT-I CD8^+^ T cells, it augmented cross-presentation of pulsed full-length ovalbumin, confirming the requirement of intracellular events for IRE1α activation. Of note, G9668 did not alter MHC-II-restricted antigen presentation to OT-II CD4^+^ T cells, nor did it affect co-stimulation of naïve CD8^+^ or CD4^+^ T cells. Enhancement by G9668 was not limited to the OT-I model, as it also applied to cross-presentation of CT26-derived antigens to splenic CD8^+^ T cells from CT26 tumor-bearing mice. Thus, the engagement of IRE1α during antigen processing within DCs selectively curtails MHC-I-restricted cross-presentation, providing negative feedback to modulate T cell activation.

Our investigation of how IRE1α curtails cross-presentation underscores RIDD as an important mechanism that depletes mRNAs encoding MHC-I heavy chains. This agrees in principle with the earlier observation of RIDD-mediated depletion of other cross-presentation components (Osorio et al., 2014; Tavernier et al., 2017), although the specific RIDD targets differ. Perhaps distinct mRNAs are depleted upon more subtle RIDD activation by antigen, as compared to RIDD hyper-activation by *XBP1* KO. Our analysis revealed the presence of consensus stem-loop endomotifs within each of the three mouse MHC-I heavy-chain mRNAs, *i.e.*, H-2K, H-2D and H-2L. We verified the cleavage of all three transcripts by the phosphorylated kinase-endoribonuclease module IRE1α *in vitro*, as well as of H-2K mRNA in antigen-pulsed BMDCs. Moreover, our scRNAseq analysis demonstrated that G9668 treatment elevated H-2K and H-2D transcript levels in tumor-associated DCs, providing further validation of this mechanism in the tumor microenvironment. Of note, multiple human HLA-I A, B and C heavy-chains also contain consensus stem-loop RIDD endomotifs (not shown). Nevertheless, we have recently observed that human IRE1α can also perform more promiscuous Rnase activity, termed RIDDLE for RIDD lacking endomotif (Le Thomas et al., submitted), which might account for divergence of IRE1α RNase targets in different settings.

In three syngeneic tumor models, systemic treatment with G9668 attenuated tumor growth more strongly than did selective *IRE1α* disruption in the malignant cells, indicating that IRE1α activity in the tumor microenvironment supports tumor growth. Flow cytometry and scRNAseq analyses demonstrated that systemic IRE1α inhibition increased MHC-I heavy chain transcript and surface-protein levels in tumor-infiltrating DCs, mirroring the *in vitro* BMDC experiments. These changes occurred in conjunction with enhanced tumor infiltration and activation of CD8^+^ T cells. Functional linkage between the enhancement of the highly specialized cDC1-mediated cross-presentation and CD8^+^ T cell engagement was strengthened by elevated MHC-I tetramer staining. Importantly, combined treatment with anti-PD-L1 antibody and G9668 in the orthotopic EMT6 TNBC model, which is only partially responsive to anti-PD-L1, led to clear tumor regression, establishing non-redundant complementarity of these two modalities.

In conclusion, our studies reveal that pulsed-antigen-derived peptides can directly engage IRE1α in cross-presenting DCs, explaining the activation of this ER-stress sensor in the absence of classical ER stress. Furthermore, by fully blocking IRE1α’s enzymatic function, we have discovered that IRE1α controls a negative feedback loop, by depleting MHC-I heavy-chain mRNAs via RIDD, to dampen cross-presentation and curtail consequent CD8^+^ T cell activation. Excitingly, disruption of this feedback by small-molecule IRE1α inhibition holds promise for cancer immunotherapy, particularly in combination with anti-PD-L1 inhibition. These findings bring important conceptual advances to seminal previous work identifying a role for XBP1s in dysregulated lipid metabolism and in function of tumor-associated DCs (Cubillos-Ruiz et al., 2015), and to studies on IRE1α disruption in immunodeficient mice (Harnoss et al., 2020; Harnoss et al., 2019; Harrington et al., 2015).

## MATERIALS & METHODS

### Cell Cultures, BMDC differentiation and Experimental Reagents

CT26, 4T1, HEK293 and EMT6 cells were originally acquired from ATCC, authenticated by analysis of short tandem repeats and tested to ensure no presence of mycoplasma within 3 mo of use. U20S cells were kindly provided by the Walter Lab of the University of California, San Francisco (UCSF). Cells were grown in RPMI1640 media supplemented with 10% fetal bovine serum (FBS) (Sigma, St. Louis, MO), 2 mM glutaMAX (Gibco, Amarillo, TX), 100 U/ml penicillin (Gibco) and 100 μg/ml streptomycin (Gibco). MEFs were obtained as previously described (Holst et al., 2007).

For purification and differentiation of BMDCs, the femur and tibia bones of C57BL/6 mice were flushed with sterile PBS and bone marrow cells were then cultured in RPMI1640 media as described above and further supplemented with 50 mM β2-mercaptoethanol (Sigma), 20 ng/ml granulocyte-macrophage colony-stimulating factor (GM-CSF) (Biolegend, San Diego, CA) and 10 ng/ml IL-4 (Biolegend) for 9 days, with media being replenished every 3 days of culturing, as described previously (Fernandez et al., 1999). Where indicated, Flt3L-BMDCs were similarly generated using media supplemented with 200 ng/ml recombinant mouse Flt3L (Peprotech, Cranbury, NJ) (Brasel, De Smedt, Smith, & Maliszewski, 2000). After 9 days of culture, BMDCs were routinely verified to be >90% CD11c^+^ MHC-II^high^ by flow cytometry analysis.

Thapsigargin (Sigma) was used at 100 nM, tunicamycin (Sigma) was used at 5 μg/ml, MG132 (Sigma) was used at 5 μM, Amiloride (Sigma) was used at 10 μM, actinomycin D (Sigma) was used at 4 μg/ml, LPS (Sigma) was used at 10 μg/ml and poly-I:C (Sigma) was used at 25 μg/ml. Ovalbumin (Sigma), endotoxin-free ovalbumin (EF-OVA) (EndoFit, InvivoGen, San Diego, CA), SIINFEKL peptide (Sigma), and human CD4-Fc fusion protein (generated in-house at Genentech) were dissolved in PBS prior to pulsing and used at indicated concentrations.

The RNase-based IRE1α inhibitor 4μ8C (Cross et al., 2012) was used at 5 μM. The kinase-based IRE1α inhibitor G9668 (Harnoss et al., 2020; Harnoss et al., 2019; Harrington et al., 2015) was used as indicated.

For antigen uptake experiments, ovalbumin and CD4-Fc fusion proteins were labelled with allophycocyanin (APC) with the Lightning-Link Labelling Kit (Abcam, Cambridge, UK).

For pulsing with tumor cell lysates, indicated cell lines were grown to confluence, suspended in sterile PBS and subjected to five freeze-thaw cycles with liquid nitrogen and heating at 37°C. Cell lysates were then normalized by BCA protein concentration measurement (Thermo-Fisher, Waltham, MA).

### *In Vitro* characterization of small-molecule IRE1α inhibitor G03089668

Potency of G9668 was analyzed in two assays of IRE1α activity, with dilutions covering a range of concentrations from 0.2 nM to 10 μM in order to determine IC_50_ values. Inhibition of RNase activity was assessed by the incubation of G9668 with IRE1α (Q470-L977) and 5^’^FAM-CAUGUCC**GC**AGC**G**CAUG-3^’^BHQ substrate. Substrate cleavage was monitored kinetically as an increase in fluorescence. Cellular activity was evaluated with the XBP1s-luciferase reporter assay in HEK293 cells stably transfected with the XBP1s-luciferse reporter construct. Briefly, cells were preincubated with G9668 for 2 hours and subsequently stimulated with Tg (100 nM) for 6 hours. IRE1α-mediated cleavage of the reporter led to luciferase expression which was detected with the addition of luciferin substrate. Kinase selectivity of G9668 against a panel of 220 kinases was measured at a concentration of 1 μM with KinomeScan^TM^ (DiscoverX, Fremont, CA). Fold selectivity was determined by IC_50_ measurement of competition by G9668 for binding of ATP to each specific kinase that showed significant inhibition by G9668 via KinomeScan^TM^.

### Generation of IRE1α KO syngeneic tumor cell lines

Individual IRE1α-specific sgRNAs were designed using a standard guide scaffold and CRISPR3. The gRNAs were cloned into pLKO_AIO_CMV_Cas9_mCherry, enabling co-expression of each sgRNA, Cas9, and an mCherry-based selection marker following transient transfection into target cells.

sgRNA target sequences used in this study:

IRE1α gRNA1: TGTTTGTCTCGACCCTGGA
IRE1α gRNA2: GAGGACGGGCTCCATCAAG
IRE1α gRNA3: GGAGGCCTGAACCAATTCT
IRE1α gRNA4: ATGTTATCGACCTCCTGAC

Transfection was with Lipofectamine 3000 (Thermo-Fisher) according to manufacturer’s protocol. At 24 hours after transfection, cells were washed once in PBS and resuspended in PBS media containing 3% BSA Fraction V. The cell suspension was then filtered through a 35 mm membrane followed by immediate FACS sorting using the mCherry selection marker. Single cell clones (n=96) were plated and grown. Clones producing colonies were tested for proper IRE1a disruption by immunoblot.

### Immunoblot analysis

Cells were lysed in PBS solution supplemented with 1X RIPA buffer (Millipore, Burlington, MA) and 2X Halt^TM^ protease-phosphatase inhibitor cocktail (Thermo-Fisher). Upon clearance, samples were analyzed by SDS-PAGE, electrotransferred to nitrocellulose membranes (Invitrogen) and blocked by 5% powdered milk in PBSt (PBS supplemented with 0.1% tween) solution. Development was conducted with ECL reagent (Thermo-Fisher) and ChemiDoc ZRS+ imager (Biorad, Hercules, CA). Antibodies used for western blot analysis include IRE1α (3294, Cell Signaling Technology, Danvers, MA), β-actin (5125, Cell Signaling), ATF6 (66563-1-Ig, Proteintech, Rosemont, IL), TAP1 (12341, Cell Signaling) CHOP (2895, Cell Signaling), ATF4 (11815, Cell Signaling), ovalbumin (P1-196, Thermo-Fisher), Myc tag (2272, Cell Signaling) and biotin (5597, Cell Signaling). IRE1α lumenal domain, XBP1s and pIRE1α antibodies were generated at Genentech. Secondary antibodies were (715-035-150, Jackson Laboratory, Bar Harbor, ME) for mouse and (711-035-152, Jackson Laboratory) for rabbit.

### Generation of TAP1 KO BMDCs

Bone marrow cells were purified from Cas9-expressing C57BL/6 mice as described above, subjected to red blood cell lysis with ACK lysis buffer and electroporated with P3 Primary Cell 4D-Nucleaofactor^TM^ X -kit (V4XP-3032, Lonza, Basel, Switzerland), as previously described (Freund et al., 2020). Once re-suspended in P3 buffer, cells were added a Cas9-ribonuceloprotein (RNP) complex (IDT) containing non-targeting or TAP1-targeting single guide RNAs (sgRNAs) (IDT). The sequences of TAP1-targeting sgRNA included: sgRNA A - GCGGCACCTCGGGAACCAAC, sgRNA B – TAACTGATAGCGAAGGCATC, sgRNA C – ACGGCCGTGCATGTGTCCCA. These sgRNA were used separately or all three combined. Bone marrow cells were then transfected with the appropriate program and grown for 9 days similarly to all other BMDC cultures.

### In vitro IRE1α LD binding assays

For experiments testing ovalbumin binding, a human IRE1α-LD-Fc fusion protein (LD-Fc) was used at indicated concentrations. LD-Fc is comprised of amino acids M1-D443 of IRE1α fused C-terminally to a linker (GRAQVTDKAARSTL) followed by the human IgG1 hinge and Fc portion. Ovalbumin (Sigma) was dissolved in sterile PBS and used in native state or denatured as indicated by incubation at 95°C or in pH 2.0 or pH 10.0 buffer for 10 min.

For plate-based binding experiments, native or heat-denatured ovalbumin at indicated concentrations was bound to a flat-bottom 96 well plate (Corning Inc., Corning, NY) with coating buffer (Biolegend), washed with PBSt, blocked with 1% BSA in PBS and subsequently incubated with IRE1α LD-Fc (10 μg/ml) in binding buffer composed of PBS with 20 mM HEPES, 100 mM KOAc and 0.2% TWEEN-20 for 2 hrs at room temperature. Plates were washed again with PBSt and incubated with an anti-human Fc HRP-conjugated antibody (ab977225, Abcam). Development was performed with TMB solution and terminated with Stop solution (Biolegend). Readings were taken with a SpectraMax M2 spectrometer (Molecular Devices, San Jose, CA).

For co-immunoprecipitation experiments, IRE1α LD-Fc (10 μg/ml) and ovalbumin (indicated concentrations) were co-incubated in binding buffer (as described above) for 2 hr and then immunoprecipitated with anti-IRE1α lumenal domain antibody (Genentech), conjugated to sepharose beads, overnight at 4°C. Beads were subsequently washed four times with lysis buffer and boiled in SDS sample buffer for 10 min. Samples were then analyzed by SDS-PAGE followed by immunoblot.

Peptide arrays were produced by the MIT Biopolymers Laboratory. The tiling arrays were composed of 18-mer peptides spanning the ovalbumin or GP70 sequences and overlapping by 3 amino acids. The arrays were incubated in methanol for 10 min and then in binding buffer (50 mM Tris pH 7, 250 mM NaCl, 10% glycerol, 2 mM DTT) for three 10 min wash cycles. The arrays were then incubated for 1 hr at room temperature with 500 nM IRE1α LD-Fc and washed again for three 10 min cycles in binding buffer to remove any unbound LD-Fc. Using a semi-dry transfer apparatus, bound IRE1α LD-Fc was transferred after washes to a PVDF membrane and detected with anti-human Fc antibody (ab977225, Abcam), ECL solution (Thermo-Fisher) and ChemiDoc ZRS+ imager (Biorad). To measure binding of IRE1α LD-Fc to each peptide, images containing developed membranes were quantified with ImageJ software (version 2.0.0). Pixel intensity was determined for all spots containing peptides, with background subtracted for spots containing no peptides. Peptides were considered to bind IRE1α LD-Fc if spot intensity was above the average of all array peptides.

For binding assays of biotin-tagged peptides, we generated a FLAG-tagged IRE1α lumenal domain (LD-FLAG) comprised of amino acids M1-D443 of IRE1α fused C-terminally to a linker (GNS) followed by a Flag tag (DYKDDDDK). LD-FLAG was incubated with synthetic N-terminal biotin-tagged peptides derived from ovalbumin in binding buffer (as described above) for 1.5 hr, cross-linked by 25 μM DSS (Thermo-Fisher) for 1 hr and subsequently incubated with 50 μM Tris (pH 7.5) for 15 min to quench cross-linking. Samples were then analyzed by SDS-PAGE followed by anti-biotin immunoblot.

### Transfection of ovalbumin-derived peptides and co-IP of Myc-tagged peptides with IRE1α

U20S or HEK293 cells were transfected with the TransIT-XL reagent (Mirus Bio, Madison, WI) with PRK-TK-Neo plasmids encoding for ovalbumin-derived peptides with a signal sequence (MGGTAARLGAVILFVVIVGLHGVRG - based on the signal sequence of Human Herpes Virus 1 Glycoprotein D, with an added lysine residue to allow signal sequence processing upon translation), a flexible linker (DLGSSG) prior to the peptide sequence, and an N-terminal Myc tag (EQKLISEE). For immunoprecipitation or real-time quantitative PCR (RT-qPCR) analysis experiments, cells were harvested 48 hr after transfection, washed twice with cold PBS, and harvested in cold PBS with protease inhibitor (Roche, Basel, Switzerland). Cells were lysed for 20 min on ice in lysis buffer (30 mM Tris, pH7.5, 150 mM NaCl, 1% Triton X-100). The lysates were cleared by centrifugation at 14,000 rpm for 10 min and then incubated with anti-Myc (Thermo-Fisher) antibody-conjugated sepharose beads overnight at 4°C. Beads were subsequently washed four times with lysis buffer and boiled in SDS sample buffer for 10 min. Samples were then analyzed by SDS-PAGE followed by immunoblot.

### Real-time quantitative PCR assay of transcript abundance

For RT-qPCR analysis, RNA was purified from BMDCs, HEK293 or U20S cells with the RNeasy Mini-kit (Qiagen, Hilden, Germany) and quantified with a NanoDrop 8000 spectrophotometer (Thermo-Fisher). Similar amounts of RNA were reverse transcribed and amplified using the Taqman^TM^ RNA-to-CT^TM^ 1-Step kit (Applied Biosystems, Waltham, MA). The following Taqman probes were used for HEK293 or U20S-derived RNA: XBP1s (Hs03929085_g1), XBP1u (Hs028565596_m1), CD59 (Hs00174141_m1), DGAT2 (Hs01045913_m1), and GAPDH (Hs02758991_g1) (Thermo-Fisher). The following Taqman probes were used for BMDC-derived RNA: H-2K (Mm01612247), CD59 (Mm00483149), DGAT2 (Mm0049536), BLOS1 (Mm00497168), RNF213 (Mm01248886), IRF7 (Mm00516793), β−2M (Mm00437762), TAP1 (Mm00443188), TAPBP (Mm00493417), ERAP1 (Mm00472842), and GAPDH (Mm99999915) (Thermo-Fisher). Assays were performed with the ViiA 7 (Applied Biosystems) system.

### *In vitro* degradation of MHC-I heavy-chain transcripts

To search for IRE1α cleavage sites within MHC-I heavy chain mRNAs, sequences were loaded unto A Plasmid Editor (APE) software and subjected to a search function for consensus GCAG locations. The location most likely to provide a stable stem-loop structure within each transcript was then chosen.

To determine cleavage by IRE1α, T7 RNA transcripts were synthesized based on cDNA templates of H-2K (#OMu17935, GenScript, Piscataway, NJ), H-2D (#MC208623, Origene, Rockville, MD) and H-2L (#MC227254, Origene). Amplification of cDNA was conducted using T7 forward primers and cDNA-based RNA was generated using HiScribe^TM^ T7 Quick High-Yield RNA Synthesis kit (New England Biolabs, Ipswich, MA). T7 RNA (1 μg) was digested at room temperature by IRE1α recombinant KR protein (1 μg) for 15 min in RNA cleavage buffer (HEPES pH 7.5, 20mM, KOAc 50 mM, MGAc 1mM, Tritox X-100 0.05%). The digestion was terminated by addition of formamide (97%) and exposed to 70°C temperature to linearize the RNA. Immediately after linearization, samples were placed on ice for 5 min and then run on a 3% agarose gel. Gels were visualized by a ChemiDoc ZRS+ imager (Biorad).

### *Ex vivo* T-cell activation and cross-presentation experiments

For *ex vivo* T-cell activation experiments, mice were euthanized and spleens were removed and mechanically disrupted with a GentleMacs tissue dissociator (Miltenyi Biotec Inc, Auburn, CA). Total spleen cells were washed with sterile PBS, counted and CD8^+^ or CD4^+^ T cells were magnetically separated with appropriate separation kits (Stemcell Technologies, Vancouver, Canada).

For CD3/CD28-mediated activation, Ultra Low-endotoxin, Azide-Free (LEAF) plate-bound anti-mouse CD3 (Biolegend) was used at 2 μg/ml and soluble anti-mouse CD28 (Biolegend) was used at 8 μg/ml. T cells were incubated for 72 hours prior to flow cytometry analysis or Cell Titer Glo (Promega) analysis.

For antigen cross-presentation assays, 2 ⋅ 10^4^ BMDCs were plated, activated with LPS (10 μg/ml) for 2 hours and pulsed with SIINFEKL (100 nM), ovalbumin, EF-OVA, or CT26 lysate (all at 250 μg/ml) overnight. BMDCs were then washed with media and 2 ⋅ 10^5^ CD8^+^ or CD4^+^ T cells were added and co-incubated for 72 hr prior to flow cytometry analysis. T cells were stained with a Celltrace Violet Cell Proliferation reagent (Thermo-Fisher) prior to introduction to the co-culture. Proliferation was then determined by loss of Celltrace Violet signal in viable T cells after co-incubation.

### Mouse strains and *in vivo* tumor growth studies

All animal procedures were approved and conformed to guidelines established by the Institutional Animal Care and Use Committee (IACUC) of Genentech and were carried out in facilities accredited by the Association for the Assessment and Accreditation of Laboratory Animal Care. In all in vivo studies, tumor size and body weight were measured twice per week. Subcutaneous and mammary fat pad tumor volumes were measured in two dimensions (length and width) using Ultra Cal-IV calipers (model 54 − 10 − 111; Fred V. Fowler Co.). The tumor volume was calculated using the following formula: tumor size (mm^3^) = (longer measurement × shorter measurement^2^) × 0.5. Tumor growth inhibition (TGI) as a percentage of vehicle was calculated as the percent difference between the daily average area under the tumor volume–time curve (AUC) of treatment and control group fits on the original untransformed scale over the same time period using the following formula: %TGI = (1-[(AUC/Day) Treatment ÷ (AUC/Day) Vehicle]) ⋅ 100.

C57BL/6 and Balb/C mice were acquired from The Jackson Laboratory or from Charles River laboratories, and MyD88 KO mice were acquired from Charles River Laboratories (Wilmington, MA). For TAP1 KO experiments, in house-generated Cas9-expressing C57BL/6 mice were used.

For CT26 and 4T1 tumor studies, mice were inoculated subcutaneously (s.c.) on the right flank. 1 ⋅ 10^5^ CT26 or 4T1 cells were counted and suspended in HBSS (Gibco) and admixed with 50% Matrigel (BD) to a final volume of 100 μl. For EMT6 tumor studies, an identical number of cells was prepared similarly and inoculated into the mammary fat pad.

For *in vivo* studies, 7 days after tumor-cell inoculation animals were randomized into groups receiving vehicle control (50% PEG400, 40% water, 10% DMSO) or G9668 (250 mg/kg) compound bidaily (BID) by oral gavage (PO). For the EMT6 combination studies, mice were randomized into four groups, with groups similarly receiving vehicle or G9668 in combination with anti-GP120 control antibody or 6E11 anti-PD-L1 antibody (both with LALAPG Fc alterations, dissolved in PBS, at 10 mg/kg intravenously (IV) for the first dose and 5 mg/kg intraperitoneally (IP) biweekly (BIW) thereafter).

### Flow cytometry

For *in vivo* experiments, tumors were excised after euthanasia, mechanically disrupted by a GentleMacs tissue dissociator (Miltenyi Biotec Inc.) and enzymatically digested by dispase (80 μg/ml, Life Technologies, Carlsbad, CA), Collagenase P (Roche, 20 μg/ml) and DNAse I (Roche, 10 μg/ml). For cytokine staining assays, cells were re-suspended in RPMI1640 growth media supplemented with T cell stimulation cocktail (4975-03, eBioscience Inc., San Diego, CA) and brefeldin A (Biolegend) and incubated at 37°C for 4 hr.

For flow cytometry analysis assays, samples were re-suspended in FACS buffer (0.5% BSA, 0.05% azide), blocked with anti-CD16/32 blocking antibodies (101302, Biolgend) for 20 min in 4°C, and then incubated with fluorescently-labelled antibodies for a further 20 min in 4°C. The following dyes were used: PI (Sigma), CellTrace Violet (Invitrogen, C34557) and Live/Dead Staining Kit (Invitrogen, L10119). The following antibodies were used: Perforin (PE, 12-9392-82, eBiosciences), ki67 (BV421, 652411), CD8 (PE-Cy7, 100722), CD4 (APC-Cy7, 100526), CD3 (APC, 100236), F4/80 (APC-Cy7, 123118), MHC Class I (FITC, 125508), MHC class II (BV605, 107639), CD11c (BV711, 117349), CD11b (APC, 101212), XCR1 (BV421, 148216), CD103 (AF488, 121408), granzyme B (FITC, 515403), CD59 (PE, 143103), PD-1 (BV605, 135220), CD44 (BV711, 103057), CD69 (PE-Cy7, 104512), IFN-γ (APC, 505810) (all from Biolegend). The Foxp3 Fix/Perm Kit was used for intracellular staining (421403, Biolegend). GP70 tetramers were generated at Genentech, as described (Vormehr et al., 2020).

Samples were read in a BD Symphony cell analyzer (BD) and data was analyzed in FlowJo Software (FlowJo 10.2, FlowJo LLC, Ashland, OR). For cell sorting, a BD FACSAria II (BD, Franklin Lakes, NJ) was utilized.

### Single cell RNA sequencing

For single cell RNA sequencing, libraries were generated using Chromium Single Cell 5′ Library & Gel Bead kit (1000006, 10X Genomics, Pleasanton, CA), from 2.5 ⋅ 10^4^ viable CD45^+^ cells sorted from 4T1 tumors.

### Statistical analysis

All values were represented as arithmetic mean ± standard deviation (SD). Statistical analysis was performed by unpaired, two-tailed *t* test or one-way ANOVA. A resulting P value < 0.05 was considered significant. All analyses were performed with the GraphPad Prism 7 software (GraphPad Software, Inc., San Diego, CA).

## AUTHOR CONTRIBUTIONS

OG designed research studies, conducted experiments and wrote the manuscript. ALT, JMH, SMH, AM and GS designed research studies and conducted experiments. SM, DL conducted experiments. LG and TW analyzed data. ZM and IM designed research studies. AA designed research studies and wrote the manuscript.

## ACKNOWLEDGEMENTS

We thank Angela Yang for assistance with design, performance and analysis of single-cell RNA sequencing of tumor leukocytes; Marie-Gabrielle Braun and Joachim Rudolph for providing G9668; Maureen Beresini for kinome selectivity analysis; David Kan and Ehud Segal for help with design and performance of *in vivo* tumor studies and treatment with anti-PD-L1 antibodies; Shannon Ruppert, Jeanne Cheung, Shiuh Luoh, Yajun Chestnut and Manmeet Singh for assistance with harvesting of tumor-bearing mice; Christine Moussion, Shannon Turley, Mable Lam, and Peter Walter for helpful discussions.

---

**Figure S1.**
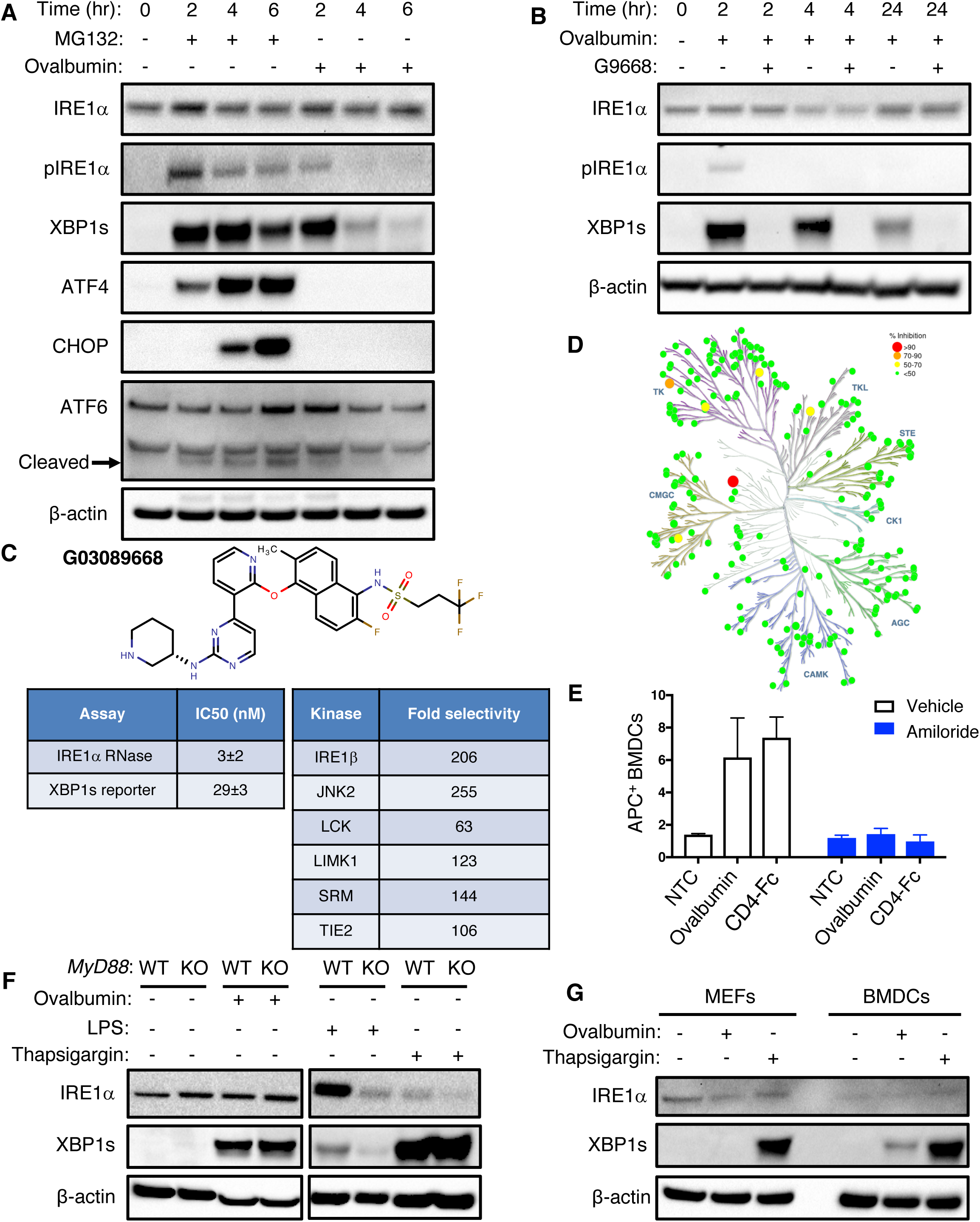
Antigen pulsing of BMDCs activates IRE1α. **(A)** BMDCs were treated with MG132 (5 μM) or ovalbumin (500 μg/ml) for indicated time periods and analyzed by IB. **(B)** BMDCs were pulsed with ovalbumin (500 μg/ml) and G9668 (3 μM) for indicated time periods and analyzed by IB. **(C)** Molecular structure, potency and kinase selectivity of G9668. **(D)** Schematic representation of G9668 interaction with 220 kinases at 1 μM. Size and color of circles are related to interaction strength. Analysis was conducted by KinomeScan^TM^ **(E)** BMDCs were pulsed for 4 hr with allophycocyanin (APC)-tagged ovalbumin or soluble CD4-Fc protein (500 μg/ml) in the absence or presence of amiloride (10 μM); uptake of protein was assayed by flow cytometry. **(F)** WT or MyD88 KO BMDCs were pulsed with ovalbumin (500 μg/ml) or treated with LPS (10 μg/ml) or Tg (100 nM) for 4 hr and analyzed by IB. **(G)** MEFs and BMDCs were pulsed with ovalbumin or treated with Tg (100 nM) for 4 hr and analyzed by IB. Images in panels **A**, **B**, **F** and **G** represent at least two similar experiments; panel **E** bar graphs represent mean ± SD from three independent biological repeats.

**Figure S2.**
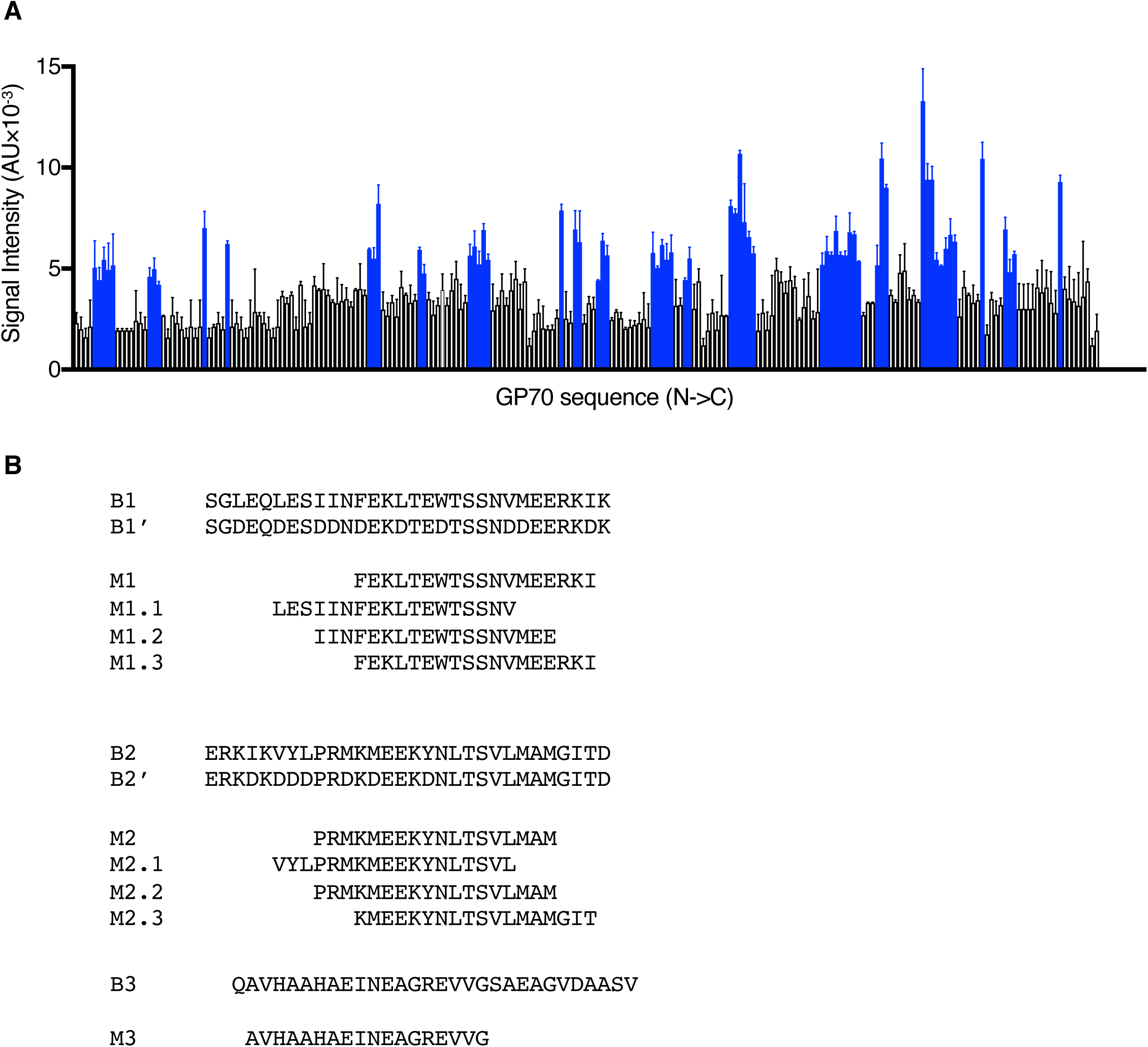
Antigen-derived peptides can directly engage IRE1α. **(A)** A tiled 18 aa-long peptide array spanning GP70 was incubated with IRE1α LD-Fc (500 nM) followed by colorimetric detection with an HRP-conjugated anti-human Fc antibody. **(B)** Sequences of biotin-tagged peptides (labeled B) used in Figure 1D; Myc-tagged signal peptides (labeled M) used in Fig. 2E; and Myc-tagged peptides (Labeled M) used in Figure 2F. Bar graphs in panel **A** represents mean ± SD from three independent technical repeats.

**Figure S3.**
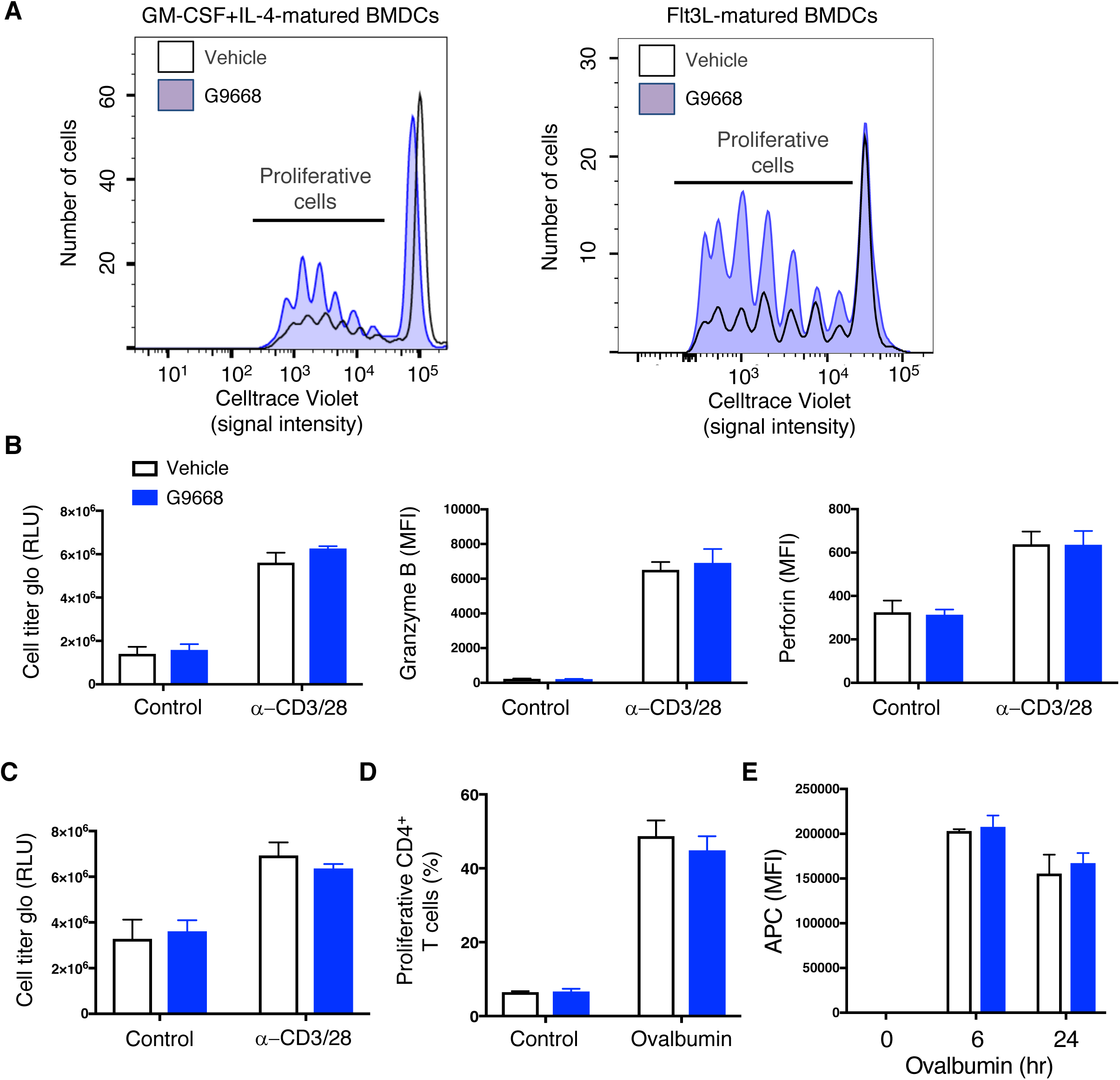
IRE1α inhibition does not affect direct TCR-mediated activation of CD4^+^ and CD8^+^ T cells; MHC-II-restricted antigen presentation and soluble antigen uptake by BMDCs. **(A)** BMDCs matured as indicated were pulsed with ovalbumin (500 μg/ml) in absence or presence of G9668 (3 μM) for 24 hr, and subsequently co-cultured with magnetically-separated CD4^+^ OT-I T cells for 72 hr, followed by flow cytometry analysis of T cell proliferation by Celltrace Violet. (**B**-**D**) Magnetically-separated CD8^+^ OT-I **(B)** or CD4^+^ OT-II **(C)** splenic T cells were activated by plate-bound anti-CD3 (8 μg/ml) and soluble anti-CD28 (2 μg/ml) antibodies in absence or presence of G9668 (3 μM) for 72 hr. Proliferation and activation were analyzed respectively by Cell Titer Glo and flow cytometry. **(D)** BMDCs were pulsed with ovalbumin (500 μg/ml) for 24 hr, with or without G9668 (3 μM) and subsequently co-cultured with magnetically-separated CD4^+^ OT-II T cells for 72 hr, followed by flow cytometry analysis of T cell proliferation by Celltrace Violet. **(E)** BMDCs were pulsed with APC-labelled ovalbumin (500 μg/ml) for indicated time and analyzed for internalization of ovalbumin by flow cytometry. **(E)** BMDCs were treated with G9668 (3 μM) for 8 hr and mRNA levels of indicated genes associated with cross-presentation were analyzed by RT-qPCR. Bar graphs in panels **B**-**E** represent mean ± SD from three independent biological repeats.

**Figure S4.**
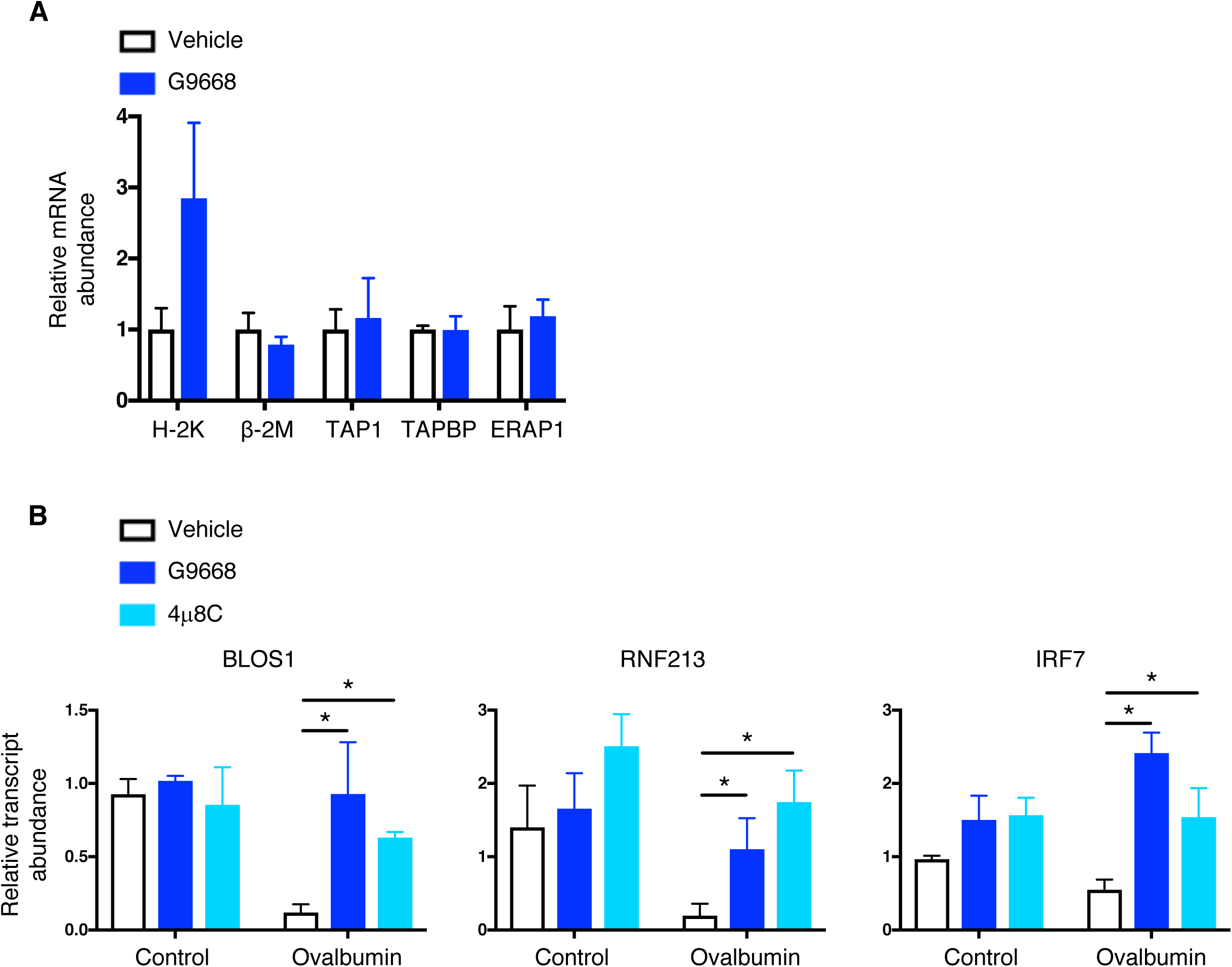
IRE1α RIDD activity specifically targets MHC-I heavy-chain transcripts. **(A)** BMDCs were pulsed with ovalbumin (500 μg/ml) for 8 hr in the absence or presence of G9668 (3 μM), followed by real time RTqPCR measurements of the indicated RIDD substrates. **(B)** BMDCs were treated with actinomycin D (2 μg/ml)) and pulsed with ovalbumin (500 μg/ml) for 8 hr combined with DMSO or G9668 (3 μM) or 4μ8C (1 μg/ml), followed by real time RTqPCR measurements of the indicated RIDD substrates. Analysis was performed using unpaired, two-tailed *t* test, * P ≤ 0.05. Bar graphs in all panels represent mean ± SD from three independent technical repeats.

**Figure S5.**
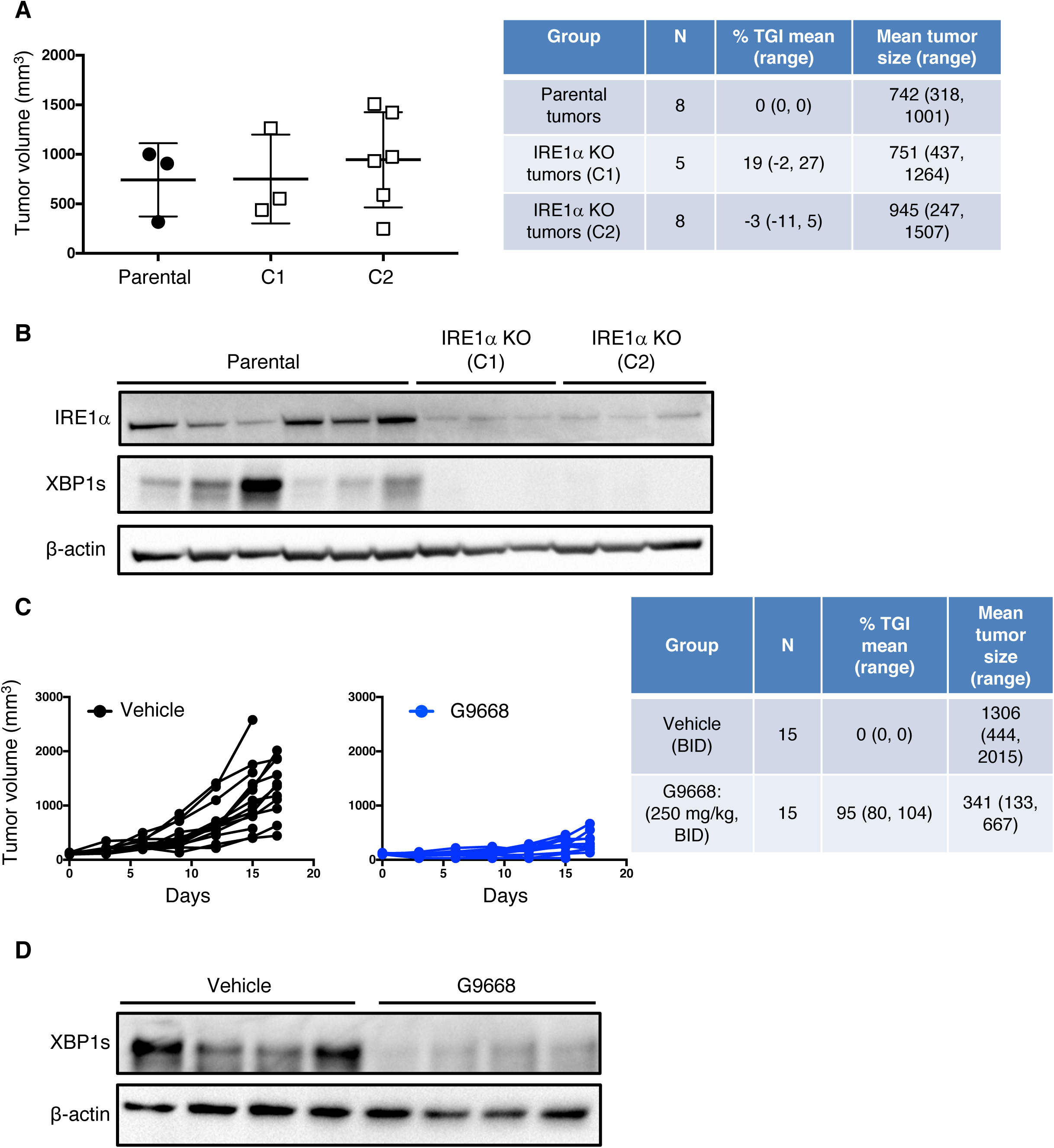
IRE1α inhibition attenuates CT26 tumor growth. **(A, B)** Animals were inoculated s.c. with parental or IRE1α KO CT26 cells and tumor growth was measured over 27 days. **(A)** Final day tumor measurements, Scatter plots represent mean ± SD. **(B)** IB analysis of IRE1α expression and activation. **(C, D)** Mice were inoculated s.c. with CT26 cells, grouped out 7 days afterwards and treated with vehicle or G9668 (250 mg/kg, BID). **(C)** Growth trajectories of CT26 tumors in individual vehicle- and G9668-treated animals over 17 days and IB analysis of total tumor lysates **(D)** are depicted. Number of animals included in each study is noted in corresponding tables.

**Figure S6.**
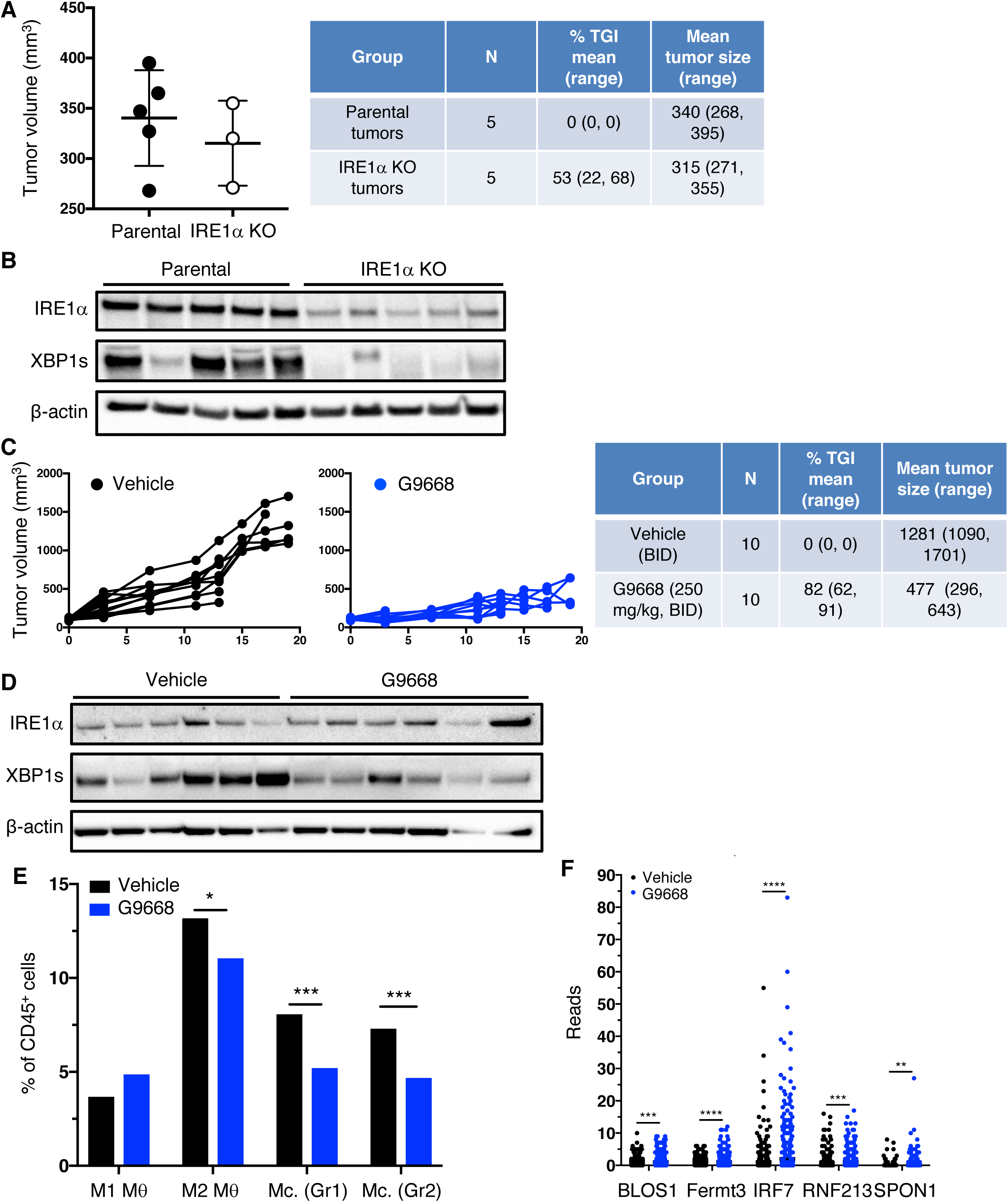
IRE1α inhibition attenuates 4T1 tumor growth. **(A-B)** Animals were inoculated s.c. with parental or IRE1α KO 4T1 cells and tumor growth was monitored over 25 days, with final measurements **(A)** and IB analysis of total tumor lysates **(B)** presented. **(C-E)** Mice were inoculated s.c. with 4T1 cells, grouped out 7 days afterwards and treated with vehicle or G9668 (250 mg/kg, BID). **(C)** Tumor growth in individual animals was measured over 19 days, and **(D)** IRE1α expression and activation were analyzed by IB. **(E, F)** Mice were treated with vehicle or G9668 for 6 days and tumors were then analyzed. **(E)** Transcript levels of indicated genes characterized as RIDD targets in tumor-infiltrating DCs. **(F)** Relative abundance of group 1 (Hcar) and group 2 (Hilpda) tumor-infiltrating monocytes (Mc.) and M1- or M2-polarized macrophages in vehicle- and G9668- treated animals, analyzed by single-cell RNA sequencing data. *P ≤ 0.05, **P ≤ 0.01 ***P ≤ 0.001, ****P ≤ 0.0001. Number of animals included in each study is noted in corresponding tables. Scatter plots in panels **A**, **E** and **F** represent mean ± SD.

**Figure S7.**
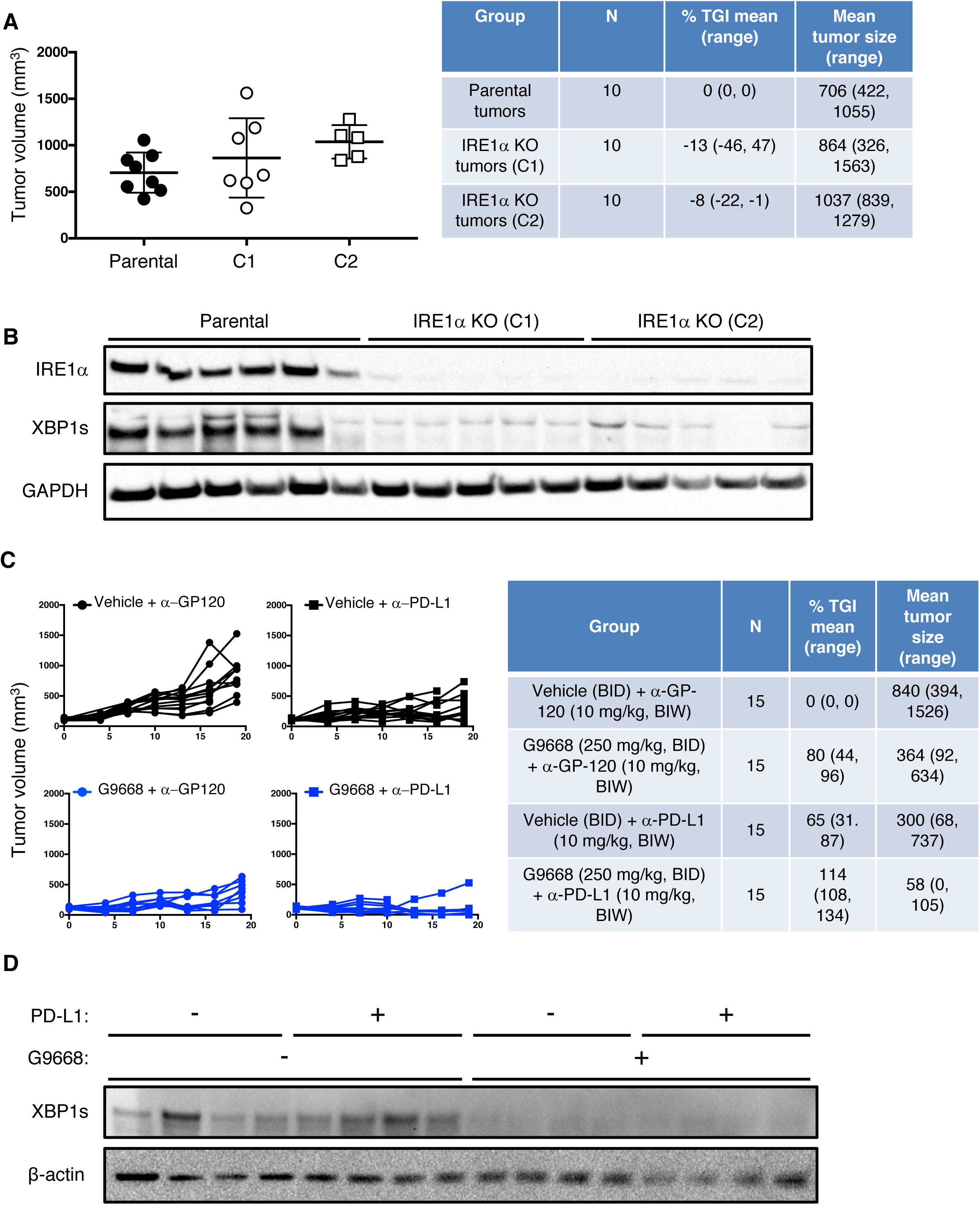
IRE1α inhibition attenuates EMT6 tumor growth and synergizes with anti-PD-L1 antibody. **(A, B)** Mice were inoculated orthotopically with WT or IRE1α KO EMT6 cells and tumor growth was measured over 24 days, with final tumor measurements (scatter plots represent mean ± SD) **(A)** and IB analysis of total tumor lysates **(B)** presented. **(C, D)** Mice were inoculated with WT or IRE1α KO EMT6 cells, grouped out 7 days afterwards and treated with vehicle, G9668 (250 mg/kg, BID), anti-PD-L1 antibody (10 mg/kg at first dose, 5 mg/kg BIW thereafter), or the combination. **(C)** Tumor growth trajectories were measured over 19 days. (**D)** IRE1α activation was analyzed by IB. Number of animals included in each study is noted in corresponding tables.

